# Adaptation to overflow metabolism by mutations that impair tRNA modification in experimentally evolved bacteria

**DOI:** 10.1101/2022.10.28.514323

**Authors:** Marc Muraski, Emil M. Nilsson, Melissa J Fritz, Anthony R. Richardson, Rebecca W. Alexander, Vaughn S. Cooper

## Abstract

When microbes grow in foreign nutritional environments, selection may enrich mutations in unexpected pathways connecting growth and homeostasis. An evolution experiment designed to identify beneficial mutations in *Burkholderia cenocepacia* captured six independent nonsynonymous substitutions in the essential gene *tilS*, which modifies tRNA^Ile2^ by adding a lysine to the anticodon for faithful AUA recognition. Further, five additional mutants acquired mutations in tRNA^Ile2^, which strongly suggests that disrupting the TilS:tRNA^Ile2^ interaction was subject to strong positive selection. Mutated TilS incurred greatly reduced enzymatic function but retained capacity for tRNA^Ile2^ binding. Yet both mutant sets outcompeted wild-type by decreasing lag phase duration by ∼3.5 hours. We hypothesized that lysine demand could underlie fitness in the experimental conditions. As predicted, supplemental lysine complemented the ancestral fitness deficit, but so did additions of several other amino acids. Mutant fitness advantages were also specific to rapid growth on galactose using oxidative overflow metabolism that generates redox imbalance, not resources favoring more balanced metabolism. Remarkably, 13 *tilS* mutations also evolved in the Long-Term Evolution Experiment with *E. coli*, including four fixed mutations. These results suggest that TilS or unknown binding partners contribute to improved growth under conditions of rapid sugar oxidation at the predicted expense of translational accuracy.

## Introduction

Evolution experiments coupled with whole genome sequencing harness the nearly deterministic effect of selection in large populations to capture beneficial mutants [1]. Any mutation that rises quickly to high frequency (e.g. within 50-500 generations) is essentially guaranteed to be adaptive or part of an adaptive genotype [2, 3]. Even though beneficial mutations are relatively rare, in bacterial populations numbering 10^7^ cells or more, numerous beneficial mutants often arise and co-occur. Under these cases, mutations of smaller benefit are likely lost because they are outcompeted by co-occurring mutants of greater benefit, which causes mutants captured first to be among the most beneficial. Surprisingly, these experiments have not captured the diversity of genetic targets that one might anticipate from a genome-wide screen of beneficial variation. Rather, experimental evolution of replicate microbial populations has revealed surprising parallelism at the gene level for early adaptations [4–6]. More surprising, the identity of repeatedly mutated genes is often unexpected, and they contribute to pathways unanticipated to play an essential role in microbial fitness. For example, in the famous Long-Term Evolution Experiment led by Richard Lenski in which *E. coli* populations were grown in a simple glucose solution for >70,000 generations [7], the first few beneficial mutations that swept to fixation in each population included a deletion of genes involved in ribose catabolism and mutations in a regulator of the stringent response, *spoT* [8, 9]. Neither of these gene targets were predicted causes of adaptation *a priori*, and 20 years after their discovery, exactly how these mutations are adaptive remains incompletely understood. Yet these and other mutants have inspired numerous subsequent studies that have improved models of how *E. coli* metabolism is regulated for optimal growth [10].

Previously, we developed a marker-deflection assay [11, 12] to capture the first beneficial mutations rising to high frequency in experimentally evolving populations of the generalist environmental bacterium *Burkholderia cenocepacia* [13]. This project was motivated by the goal of comparing the spectrum of captured beneficial mutations in a metabolically versatile microbe like *B. cenocepacia* with those identified in a domesticated laboratory strain of *E. coli* using similar methods [14]. We hypothesized that a broader range of adaptations would be captured in the environmental microbe than in the laboratory strain, but our results quickly rejected this hypothesis and led us to focus on the highly unusual beneficial mutations identified by whole-genome sequencing (WGS).

We report strong positive selection on six, nonsynonymous single nucleotide substitutions in the *tilS* gene in *B. cenocepacia* (strain HI2424). This gene encodes tRNA^Ile^ lysidine synthetase (TilS) and is considered essential, meaning that deletion mutants are inviable [15–17]. TilS modifies the tRNA isoleucine acceptor (tRNA^Ile2^) to decode the minor isoleucine codon AUA [18]. Most bacterial species encode a tRNA^Ile2^ with a CAU anticodon, which it shares with tRNA^Met^, instead of a Watson-Crick complementary UAU anticodon. To avoid mistranslation, post-transcriptional modification by TilS converts the tRNA^Ile2^ wobble nucleotide C34 to lysidine (L) by adding lysine to this cytosine, which shifts base pairing specificity [15]. Similarly, the presence of lysidine in tRNA^Ile2^ shifts aminoacyl-tRNA synthetase (AARS) preference from methionyl-tRNA synthetase (MetRS) to isoleucyl-tRNA synthetase (IleRS) [15, 19]. Because enzymatic modification of tRNA^Ile2^ is essential to maintain translational fidelity, the discovery of multiple mutations in *tilS* associated with adaptations in laboratory culture is altogether unexpected.

Further, we identified five additional mutants from these same evolution experiments in the tRNA^Ile2^ that is modified by TilS. Together, these mutants imply strong selection to alter the interaction between TilS and tRNA^Ile2^ to improve *B. cenocepacia* fitness in a defined laboratory medium. Here, we propose and evaluate possible hypotheses for these results that could connect translational fidelity to improved growth and metabolism with galactose as sole carbon source. We discovered that rapid growth in a minimal galactose medium with insufficient buffering capacity caused overflow metabolism, acidification, and redox imbalance, in which *tilS* and tRNA^ile2^ mutants were favored. These studies demonstrate new connections between central metabolism and translational fidelity in which bacteria may sacrifice protein accuracy by altering essential gene products when metabolic constraints demand.

## Materials and Methods

A summary of our approach follows. More detailed methods are described in Supplemental Text.

### Bacterial strains and culture conditions

*B. cenocepacia* HI2424, a soil isolate of a globally distributed clone causing infections in persons with cystic fibrosis and other immunological disorders, was the wild-type (WT) ancestor used in all experiments and is naïve to laboratory conditions [20]. A lacZ+ mutant was created by Tn7 insertion that is neutral for fitness but distinguishable on X-gal plates [21]. Selection experiments were conducted in a modified version of M9 minimal media with 1% galactose (GMM) added as the sole carbon source. Equal fractions of HI2424^lac^ and HI2424^lac-^ genotypes were added to 5 mL GMM in test tubes grown on a roller drum and propagated by 1:100 dilutions for 6 days and 1:10,000 dilutions for up to 6 additional days. Every 72 hours, a 10^−4^–10^−5^ dilution of each culture was counted on X-gal plates to determine marker frequency; when it diverged beyond 3:1 or after 12 days (∼120 generations), single clones were picked from both winning and losing marker types and compared with the ancestor for altered fitness. Clones found to be more fit (selection rate constant r > 0 [22] during 24h of direct competition with the oppositely marked WT strain) were genotyped by whole genome sequencing as described [5].

### Fitness assays

Fitness was measured by growth curves in GMM mimicking selective conditions, except when supplements were added, e.g. 0.60 mM L-arginine, 0.10 mM L-aspartate, 0.40 mM L-lysine, 0.10 mM L-methionine, 0.20 mM L-phenylalanine, 0.05 mM L-tryptophan, or 0.18 mM iron(III) chloride hexahydrate; the modified M9 with 0.1% galactose or 1% mannose; or the standard M9 minimal medium recipe [23] with 1% galactose.

### TilS enzyme activity

His-tagged wild-type and mutant TilS proteins were purified from *E. coli* expression vectors. Lysidinylation activity was monitored as previously described [24, 25]. Efficiency of tRNA^Ile2^ binding was quantified by electrophoretic mobility shift assays and by northern blot. Cellular lysidine levels were determined by LC-MS as described [26]. RNA-sequencing of mutants and WT were conducted as described previously [27] with modifications described in Supplemental Text.

## Results

### Strong genetic parallelism among adaptive mutations

We used a marker-deflection assay to identify early beneficial mutants of a clone of *B. cenocepacia* str. HI2424 (wild-type, or WT), initially isolated from soil [20] when growing in minimal media containing galactose as the sole carbon source. Equal fractions of otherwise identical lac^+^ and lac^−^ strains were inoculated in replicate tubes and propagated daily by 1:100 dilution for 6 days followed by 1:10,000 dilution for 6 additional days. Dilution levels were lower at the start to avoid extinction of the ancestral population, which had a long lag phase. If the initial 1:1 ratio of markers deviated to 3:1 in either direction, or after these 12 days (∼102 generations), the experiment was stopped, and clones of both marker types were picked as putative “winner” and “loser” genotypes (Figure 1A). These putative mutants were competed against the ancestral genotype of the opposite marker over 24h in the same experimental conditions, and if they outcompeted the ancestor (relative fitness r > 0, Supplementary Table 3) their genomes were analyzed by WGS to identify the causative adaptive mutations.

**Figure 1:**
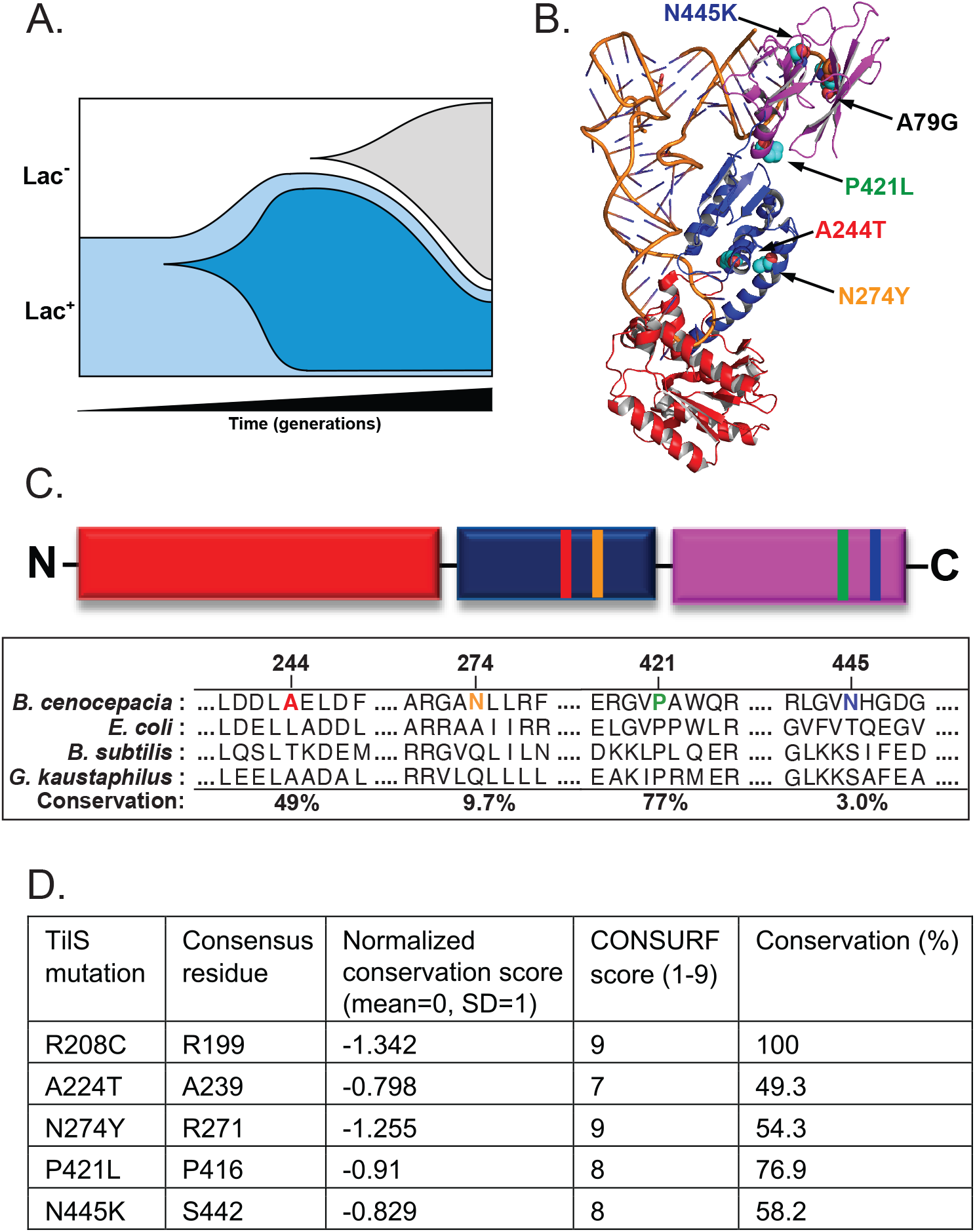
Identification of *B. cenocepacia* TilS and tRNA^Ile2^ mutants that increased fitness in minimal medium containing 1% galactose. **A**. Schematic of marker deflection assay, where beneficial mutations can arise in either the Lac^+^ or Lac^−^ ancestor population. **B**. Structure of TilS:tRNA^Ile2^ complex indicating location of single nucleotide substitutions in both partners at locations distal from the enzyme active site. Red, catalytic domain; blue, linker domain; purple, acceptor stem binding domain. Shown is the *G. kaustophilus* TilS:*B. subtilis* tRNA^Ile2^ structure, PDB 3A2K [24]. **C**. Schematic of TilS domain organization with mutation locations noted using the same color scheme as panel B, and degree of conservation of mutated residues among 150 phylogenetically diverse orthologs ranging from 35-95% overall identity using CONSURF [28]. A static CONSURF analysis of TilS can be found for PDB 3A2K at https://consurfdb.tau.ac.il/main_output.php?pdb_ID=3A2K&view_chain=A&unique_chain=3A2K A. **D**. Evolutionary conservation of mutated TilS residues. Normalized conservation scores are set with mean of 0 and one standard deviation equal to +1 or −1. CONSURF scores report similar results on a scale of 1 (least conserved) to 9 (most conserved).

In total, 19 putatively beneficial mutants were selected from three independent marker-deflection assays with the WT strain of HI2424 (Supplementary Table 4). Of these sequenced genotypes, 11 acquired single nucleotide mutations, seven had 2-4 mutations each, and two had no detectable mutation and were not considered further (Supplementary Table 4). Strong parallelism in the altered genes was evident: six genotypes acquired nonsynonymous mutations in *tilS*, encoding tRNA isoleucine lysidine synthetase (TilS); four genotypes were mutated in *ppc*, encoding phosphoenolpyruvate carboxylase; and one SNP occurred in tRNA^Ile2^, the substrate of TilS. This report focuses on the mutations in *tilS* and tRNA^Ile2^ (Figure 1B); a study of the *ppc* mutants will appear separately. Four of the six *tilS* mutants had only a single nucleotide substitution in that gene and no other, demonstrating that these mutations are the definitive genetic cause of the fitness advantage in minimal galactose medium.

We repeated this same design using a pre-adapted clone of *B. cenocepacia* that had been propagated in a similar medium but also selected for growth on plastic beads [29], again using marker-deflection versus an isogenic ac^−^ mutant to capture new mutants. In total, 15 new mutants were isolated that included six with a single mutation and nine with 2-4 new mutations (Supplementary Table 5). Notably, four mutants acquired substitutions in tRNA^Ile2^ and nine acquired mutations in *ppc*, providing further evidence of genetic parallelism of adaptations. Because isolating the phenotypic contributions of these parallel mutations was complicated by the seven mutations this ancestor had previously acquired in an earlier experiment [29], we did not analyze these genotypes further. However, the parallel evolution of six nonsynonymous mutations in the essential *tilS* gene as well as five mutations in its cognate tRNA^Ile2^ gene supports the inference that selection acted to disrupt the interaction between these gene products. These genotypes became the experimental focus of this study.

To evaluate the evolutionary conservation of mutated TilS residues and whether they may be tolerant of variation, we aligned TilS amino acid sequences to the co-crystal structure of *Geobacillus kaustaphilus* TilS and *Bacillus subtilis* tRNA^Ile2^ (PDB 3A2K) [24] using CONSURF [28]. The positions of mutated residues are highly conserved and all are among the top 25% most conserved residues among 150 phylogenetically diverse homologs (Figure 1C, 1D). We note that while mutations evolved at highly conserved positions, the parental BcTilS residue does not match the consensus residue in all cases. For example, N274 aligns with the consensus R271; this residue is only Asn in 9.7 % of the sequences. Likewise, N445K aligns with consensus S442 and is only Asn in 3.0% of the sequences. Of the six TilS mutants identified in the WT background (R208C, N274Y, P421L, N445K, and A244T found twice), all but R208C occurred alone, meaning that these single substitutions are the sole cause of the phenotypes reported here. Because R208C was not isogenic, this mutant was not studied further, although notably this residue is strictly (100%) conserved.

### Mutations in *tilS* and *tRNA*^*Ile2*^ enhance growth in the absence of certain amino acids

We hypothesized that the parallel evolution of mutations in *tilS* and its cognate tRNA^Ile2^ resulted from selection to remedy some metabolic inefficiency tied to amino acid availability, given that lysine is a substrate of TilS. We therefore compared growth kinetics of *tilS* and tRNA^Ile2^ genotypes with WT in the galactose minimal media (GMM) in which they originally evolved and in GMM containing various nutritional supplements. As expected, fitness, defined as area under the growth curve (AUC), was significantly greater for *tilS* and tRNA^Ile2^ mutants in GMM (Figure 2). It is notable that fitness advantages of all mutants were statistically similar in most assays, suggesting that either disrupting the enzyme (TilS) or the substrate (tRNA^Ile2^) is equivalently beneficial (Supplementary Figure 1). Supplementation with lysine or with aspartate, arginine, or isoleucine, each of which can be efficiently converted to lysine, complemented the fitness defect of WT and eliminated significant differences between genotypes (Figure 2 and Supplementary Figure 1). Consistent with this model that lysine deficiency favored these mutants, supplementation with amino acids that are metabolically distant from lysine (methionine, tryptophan) had no effect on fitness difference, although phenylalanine, which is not tied to lysine pathways, did complement WT fitness and improved growth of all mutants (Figure 2 and Supplementary Figure 1). The results suggest that scarcity of certain free amino acids including lysine contributed to selection of the mutants.

**Figure 2.**
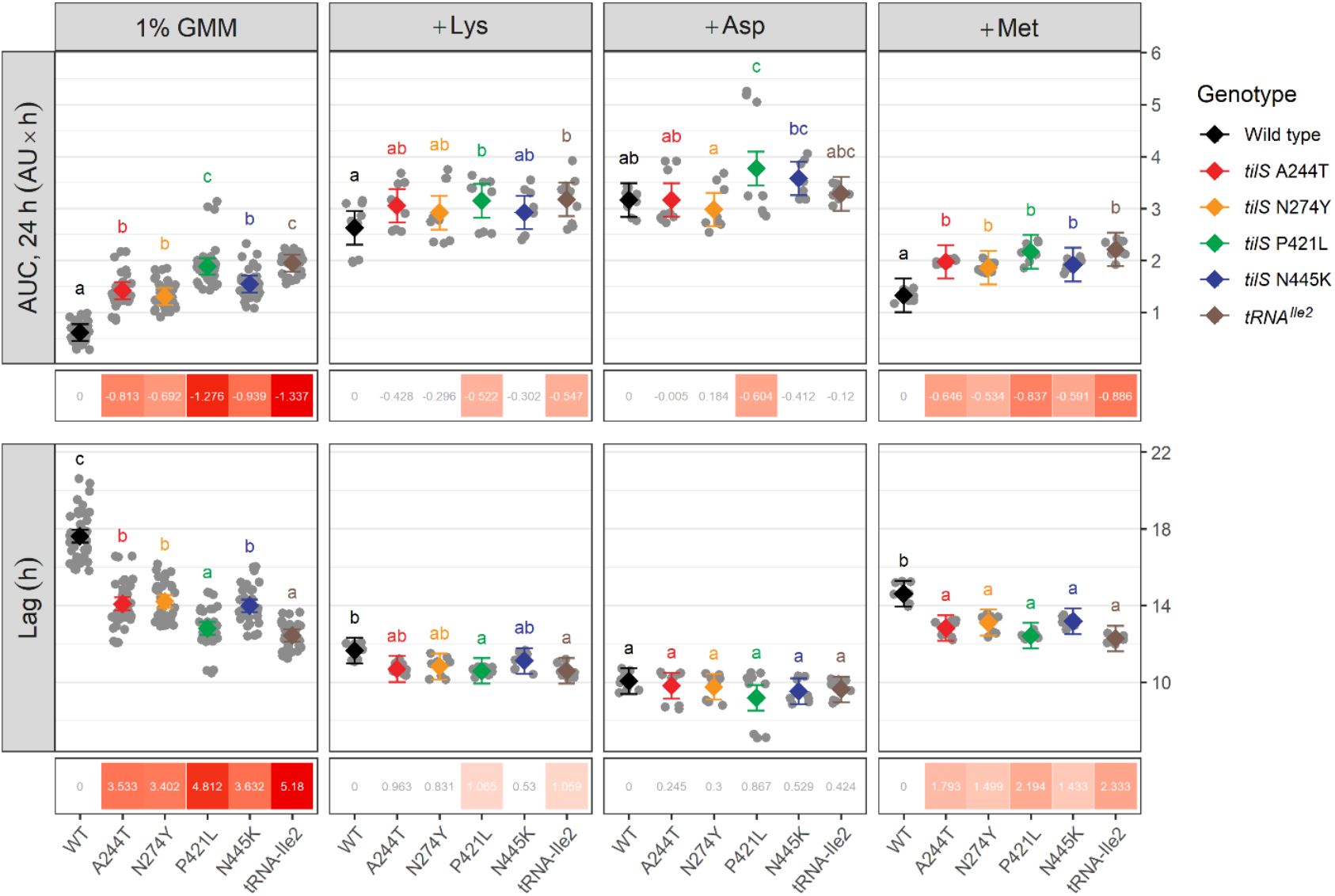
Fitness components of WT and *tilS* or *tRNA*^*Ile2*^ mutants of *B. cenocepacia*. Increased fitness of *tilS* mutants (area under the curve, AUC, top row) results from reduced lag phase (Lag, bottom row) in galactose minimal medium (GMM), but the fitness deficit of wild-type (WT) is complemented by added lysine (Lys) or aspartate (Asp) but not methionine (Met). Means and confidence intervals are in black (WT) or color (mutants), with individual observations in grey. Letters distinguish statistically significant groupings analyzed by pairwise means testing after two-way ANOVA, p<0.05. Quantitative differences in fitness measures (WT-mutant) are shown in boxes at bottom, and significant differences are shaded by magnitude of difference. AUC = area under the curve. AU = absorbance units at OD_600_.

Fitness benefits of mutants were further amplified over 36 hours of incubation relative to the 24-hour interval of serial transfer in the selection experiment. The growth curve dynamics (Figure 3A) indicate why: cultures did not reach stationary phase until at least 36h. These dynamics also show that fitness benefits of mutations arise mostly from the ability to emerge from lag phase approximately three hours earlier than WT (Figure 3A). On the contrary, the maximum growth rates of the mutants are indistinguishable from their ancestor regardless of supplementation (Supplementary Figure 2).

**Figure 3.**
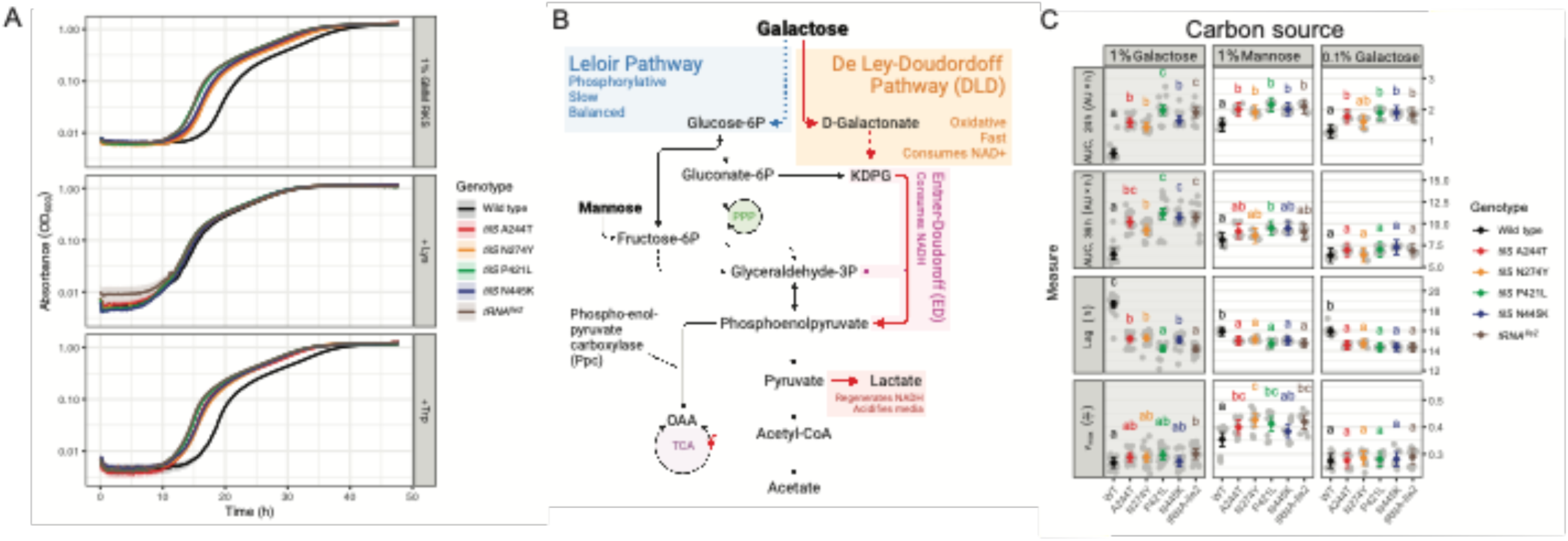
Fitness advantage of *tilS* and *tRNA*^*Ile2*^ mutants is specific to imbalanced oxidative growth on substrates like galactose and in the absence of amino acids like lysine. **A**. Lysine supplementation eliminates delayed lag of *B. cenocepacia* WT relative to *tilS* and *tRNA*^*Ile2*^ mutants. Added lysine (middle panel), but not tryptophan (bottom) to the selection media (unaltered, top) complements the WT fitness defect relative to *tilS* mutants by reducing lag phase duration. Several other substrates eliminate fitness differences between WT and mutants (Supplementary Figure 1). **B**. Identity and concentration of the carbon source dictates relative fitness advantage of *tilS* mutants. Galactose can be metabolized via either the slower, redox-balanced Leloir pathway, or the faster combination of De Ley-Doudoroff (DLD) and Entner-Doudoroff (ED) pathways that causes redox imbalance and media acidification. Note that phosphoenolpyruvate carboxylase (Ppc) provides a shunt past pyruvate and avoids lactate production. **C**. Fitness differences are greatest in 1% galactose (Gal), followed by 0.1% galactose, and least in 1% mannose (Man). Means and confidence intervals are in black (WT) and color (mutants), with individual observations in grey. Different letters distinguish statistically significant groupings determined by pairwise means testing after two-way ANOVA, p<0.05. Grey panel background denotes GMM selective medium. AUC = area under the curve. Lag and maximum growth rate (*V*_*max*_) inferred from growth curves as described in methods. AU = absorbance units at OD600.

### Fitness is dependent on carbon source and concentration

The evolved mutants were isolated in selective minimal media with excess galactose as the only available carbon source. To test whether their fitness advantages were specific to this carbon source or its concentration, we substituted galactose with mannose or varied sugar concentrations. Mannose enters metabolism through the Leloir pathway via a fructose intermediate, whereas galactose enters through either the Leloir or the De Ley-Doudoroff (DLD) pathway, followed by the redox-imbalanced Entner-Doudoroff (ED) pathway (Figure 3B). Given the high galactose concentration, we hypothesized that this sugar was predominantly metabolized through the DLD pathway via a galactonate intermediate, as the Leloir pathway is quickly overwhelmed [30]. As predicted, substituting mannose for galactose greatly reduced the fitness advantage of the mutants (Figure 3C). Furthermore, when galactose concentration was reduced from 1% to 0.1%, the growth curves more closely resembled that in 1% mannose. These results suggest that mutant fitness advantages depend on both amino acid scarcity and the primary pathway for substrate uptake and metabolism. Following research demonstrating that rapid growth on glucose by *Burkholderia* also relies upon the DLD-ED pathways [30], we also measured fitness in both 1% and 0.1% glucose. As predicted, all mutants grew earlier and to a higher density than WT in both concentrations, but with a greater advantage at higher glucose levels (Supplementary Figure 3).

Another predicted outcome of galactose catabolism in *Burkholderia* is acidification of the media via the Entner-Doudoroff (ED) pathway, which induces secretion of organic acids as a byproduct to restore redox balance [30]. Rapid growth on galactose and flux through the ED pathway causes increased production of glyceraldehyde-3-phosphate and pyruvate, both of which consume NADH (Figure 3B). Pyruvate is subsequently converted into lactate, some of which is secreted and causes media acidification. In prior experiments with this strain and growth media, we noticed acidification during growth but did not recognize this metabolic process [31]. The fitness advantages of *tilS* and tRNA^Ile2^ mutants in GMM are largely explained by their reduced lag phase and hence earlier entry into exponential growth (Figure 3A, 3C), but this change in growth could also affect the rate at which the medium acidifies. We measured the relationship between cell density and media pH for representative genotypes *tilS* N274Y and WT, in both the poorly buffered GMM media used in the selection experiment and in the standard M9 buffer (CSH) with 1% galactose (Supplementary Figure 2). As expected, N274Y grows faster than WT and reduces pH below 6.5 by 24h and below 6.0 at 36h. The WT strain eventually reaches similar OD600 and pH levels but approximately 4 hours later, which translates into a large disadvantage in direct competition when the earlier-growing mutant competitor is acidifying the growth environment. On the contrary, growth in 1% mannose or 0.1% galactose does not acidify the media as quickly (Supplementary Figure 3). Because these growth curves were conducted in separate cultures, they cannot account for possible interactions between genotypes related to greater acid tolerance of mutants, suppression of the slower-growing WT, or metabolic cross-feeding. In summary, *tilS* mutants grow better on resources like galactose and glucose (Supplementary Figure 3) that are metabolized by the De Ley-Doudoroff (DLD) and Entner-Doudoroff (ED) pathways, which causes redox imbalance and media acidification. Mutants acidify the medium faster, tolerate this stress, and indirectly limit growth of late-growing competitors, but they do not alter the relationship between growth and pH in these experimental conditions.

### Mutations in *tilS* greatly reduce lysidinylation of tRNA^Ile2^

The metabolic advantages of selected *tilS* and tRNA^Ile2^ mutations suggest that altered catalytic or binding functions of TilS with its cognate tRNA are responsible for *B. cenocepacia* fitness differences. The selected nonsynonymous mutations are distant from the TilS active site (Figure 1B) but nonetheless alter a gene identified as essential for translation [16, 17, 32]. We hypothesized that the mutations would impact TilS enzymatic function and tested this by measuring activity and structure of recombinant over-expression constructs of four TilS mutants in *Escherichia coli*. The influence of each mutation on TilS structure was assessed via circular dichroism. Only the N274Y variant displayed a modestly different α-helical content in its folded state compared to wild-type TilS (Supplementary Figure 4), suggesting that most mutations had minimal effects on protein structure. [It should be noted that the N274Y variant also required lower IPTG induction to generate soluble protein.] We assayed each recombinant protein variant for its ability to catalyze lysidine synthesis using *in vitro* transcribed tRNA^Ile2^. Each mutation lowered catalytic activity compared to the wild-type enzyme, with A244T, N274Y, and P421L substitutions reducing activity by 30-fold to >100-fold (Figure 4A) and N445K mutation reducing activity by 6-fold (Supplementary Table 7). Each selected mutation therefore caused loss of function *in vitro* despite affecting residues distal to the active site: residues 244 and 274 are about 25 Å away from the active site, while residues 421 and 445 are 60 Å and 75 Å away, respectively. This observation leaves open the possibility that the diminished activity of selected TilS mutants may be due to impaired tRNA binding.

**Figure 4.**
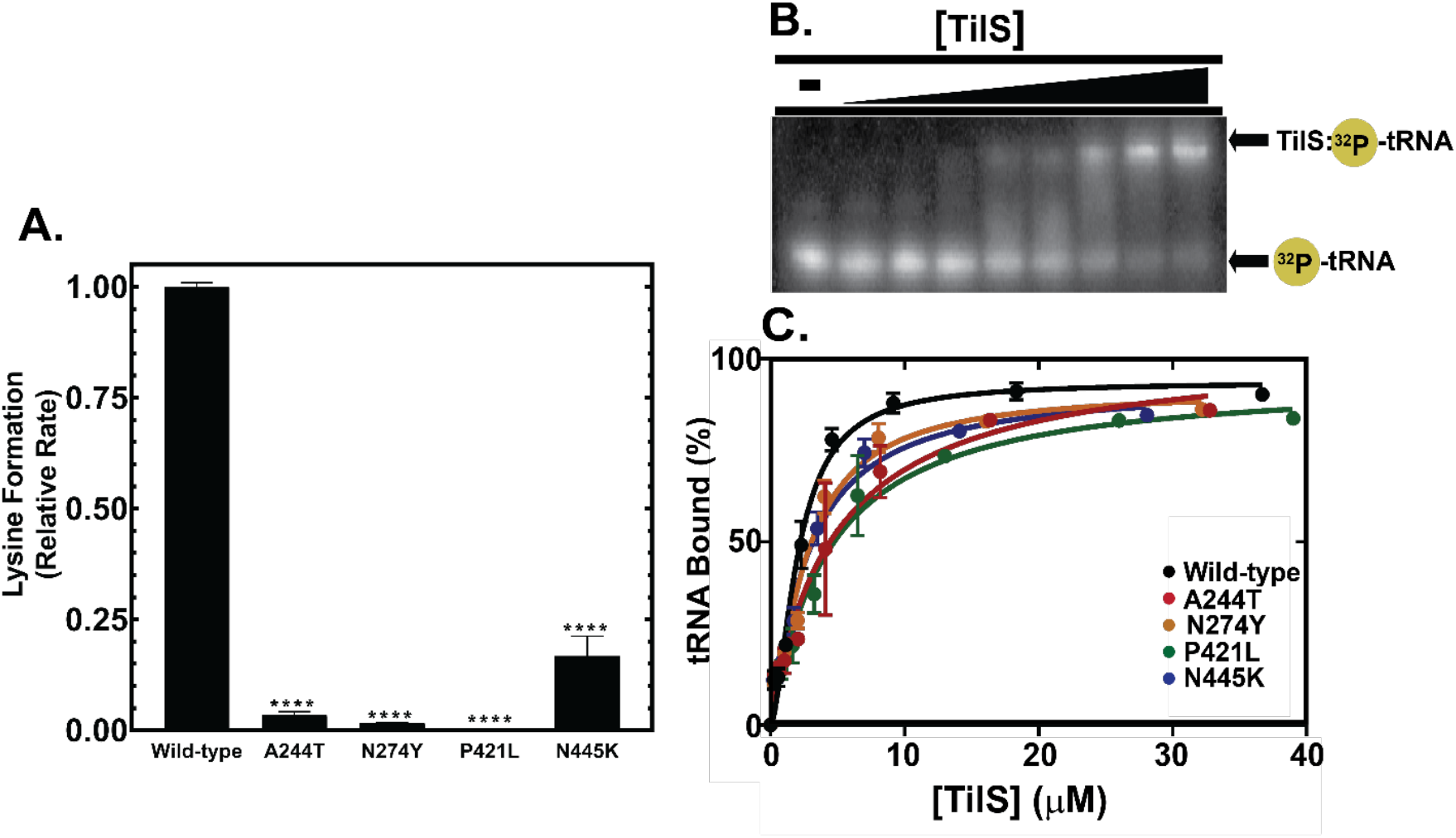
Evolved TilS mutations reduce catalysis significantly and tRNA binding modestly. **A**. Catalytic activity of *B. cenocepacia* TilS variants. Lysidinylation activity was normalized to wild-type TilS; each protein variant was determined to be significantly (p<0.05) different from wild-type using a one-way ANOVA analysis. **B**. Representative EMSA gel of TilS:tRNA^Ile2^ complex formation. ^32^P-radiolabeled tRNA^Ile2^ is separated from the ribonucleoprotein complex on a 10% native polyacrylamide gel. **C**. Graphical summary of EMSA data fit to the Hill equation in Prism v. 7.0 (GraphPad).

### Evolved TilS mutations reduce but do not eliminate tRNA^Ile2^ binding

To determine whether mutants affected the ability of TilS to complex with tRNA^Ile2^, we conducted electromobility shift assays (EMSAs) by incubating ^32^P-labeled tRNA^Ile2^ with increasing amounts of TilS variants. Mutants exhibited reduced binding affinity for tRNA^Ile2^ (Figure 4C, Supplementary Table 6) that likely contributes to the catalytic defect, but the <2-fold reduction in K_d_ seems too small to explain the observed loss of enzymatic function approaching two orders of magnitude. These results suggest that reduced catalysis by TilS variants is not simply due to an inability to recognize tRNA^Ile2^. We next asked whether mutants altered the *in vivo* pool of modified tRNA.

### tRNA^Ile2^ transcription levels are reduced in some mutants and diminish lysidine incorporation *in vivo*

One possible mechanism by which the cell can overcome diminished enzyme activity is by increasing synthesis of the enzyme’s substrate. We investigated whether the evolved mutants overproduced tRNA^Ile2^ *in vivo* by probing total *B. cenocepacia* RNA with CY5-labeled oligoDNAs targeting BctRNA^Ile2^ or BctRNA^Met^. The northern blot analysis revealed no increase in tRNA^Ile2^ levels, but rather a modest decrease in the P421L mutant and a ∼50% reduction in the A79G mutant (Figure 5A). To test whether TilS mutants indeed failed to modify cellular tRNA^Ile2^, levels of individual nucleosides were analyzed by HPLC-coupled mass spectrometry. Lysidine modification is exclusive to tRNA^Ile2^, so loss of lysidine in the tRNA pool correlates to the loss of TilS activity. Lysidine levels in the TilS mutants mimicked the catalytic trends of these transcripts *in vitro* (Figure 4A), with each of the mutant strains demonstrating reduced lysidine from the tRNA pool as compared to the WT ancestor (Figure 5C). Mutants A244T, N274Y, or N445K produced only ∼30% of the lysidine level seen in the WT ancestor, while P421L produced only 10% of WT levels (Supplementary Table 6). Thus, the catalytic phenotypes observed *in vitro* are broadly consistent with those *in vivo*, and more importantly, increased fitness during growth in minimal galactose media is tied to the loss of the only known conserved function of the TilS enzyme.

**Figure 5.**
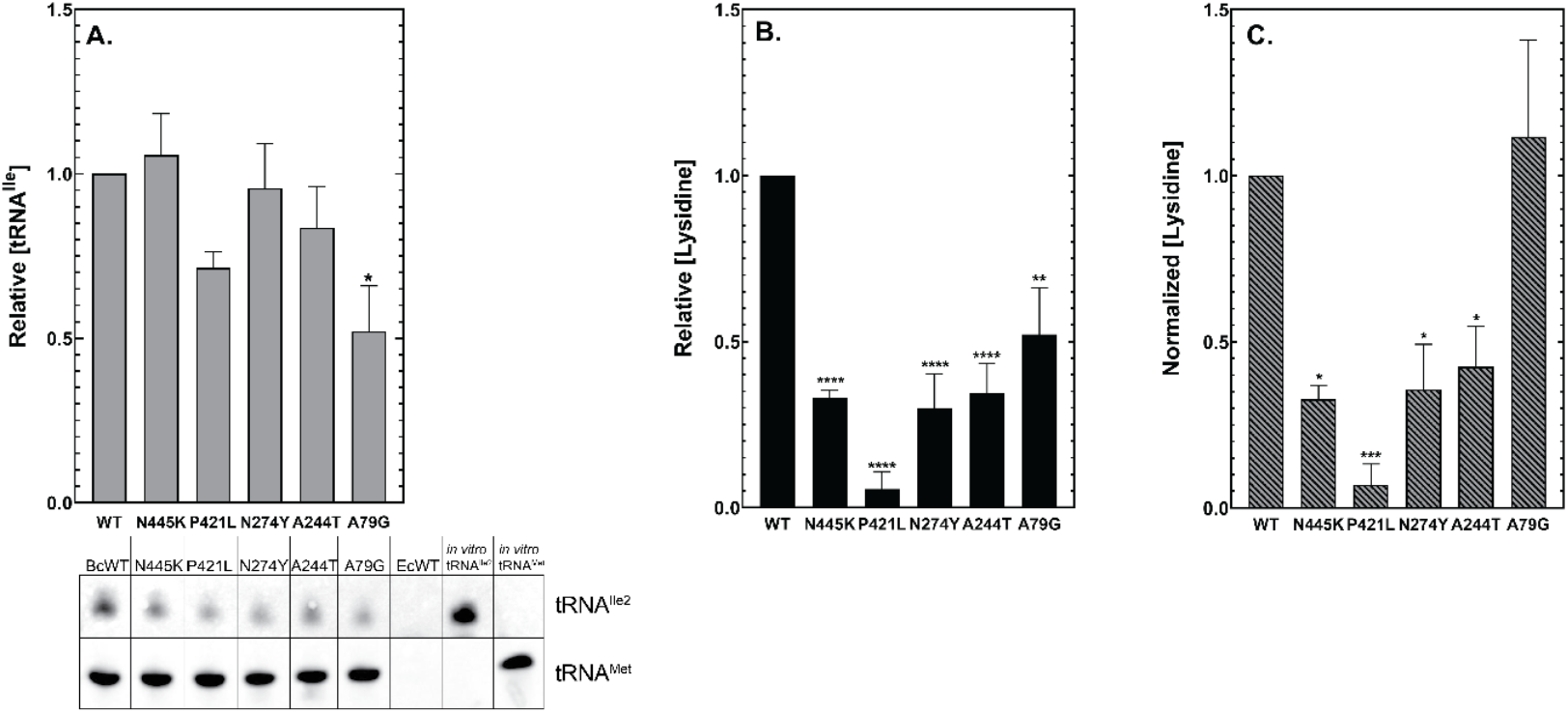
Cellular lysidine levels are decreased in TilS mutants but not in a tRNA^Ile2^ mutant. Total RNA was extracted from each strain, including tRNA^Ile2^A79G (A79G). **A**. CY5-labeled probes specific for *B. cenocepacia* tRNA^Ile2^ or tRNA^Met^ were used to determine cellular tRNA abundance as a percentage of WT by northern blot. Total *E. coli* RNA was used as a negative control to demonstrate probe specificity; *in vitro* transcribed *B. cenocepacia* tRNA^Ile2^ and tRNA^Met^ were used as positive controls. **B**. Nuclease P1-digested total RNA was analyzed by LC-MS for the presence of lysidine. **C**. Detected lysidine (panel B) was normalized to tRNA^Ile2^ levels in each sample (panel A) to adjust for differences in tRNA transcription levels between each sample and replicate; by ANOVA with post-hoc testing for five biological replicates. (* = p 0.05; ** = p 0.01; *** = p 0.001; **** = p 0.0001)

### *tilS* mutations are broadly pleiotropic and their transcriptomes indicate enhanced metabolic efficiency

We predicted that the large fitness gains of *tilS* and tRNA_Ile2_ mutants would associate with altered expression of gene sets that would suggest mechanisms of their growth advantage. Comparing mutant and WT transcriptomes was complicated by their different growth dynamics, rates of resource consumption, and media acidification (Figure 3, Supplementary Figure 3). To evaluate genotype differences more independently of these environmental feedbacks, we isolated RNA from each replicate at equivalent optical density (but different time points) for RNA-seq. Among hundreds of differently expressed genes, most outliers belong to four pathways (Figure 6). First, each mutant showed upregulated iron uptake through Fe_3+_ siderophore receptors like ornibactin (*orbA*) and regulators like the FecI sigma factor (Bcen2424_1359), which are likely required to assemble iron-sulfur proteins that are in high demand during bacterial lag phase [33]. Second, mutants upregulated the glyoxylate bypass through the iron-sulfur cluster enzyme aconitase (*acnB*), which was the transcript showing the greatest fold-increases, and isocitrate lyase (*icl*), which preserves carbon for biomass generation [34]. Two additional pathways are strongly suppressed in mutants: production of polyhydroxylalkanoates (PHAs), energy storage molecules that are typically produced in large quantities by *Burkholderia* but which consume acetyl-CoA, and production of a newly discovered antifungal secondary metabolite called fragin, whose expression demands directly oppose that of *tilS* mutants [35]. These differences are clearly illustrated in comparing N274Y with WT (Figure 6), but are representative of other mutants including *tRNA*_*Ile2*_, which shows similar effects on expression but differences of lower magnitude (Supplementary Figure 5). Together, these data suggest that un-modified tRNA_Ile2_ or unoccupied TilS generate a signal or process that increases iron acquisition and conserves carbon as acetyl-CoA for synthesis of all cellular components from the sole substrate of galactose, while reducing investment in energy storage and metabolic by-products.

**Figure 6.**
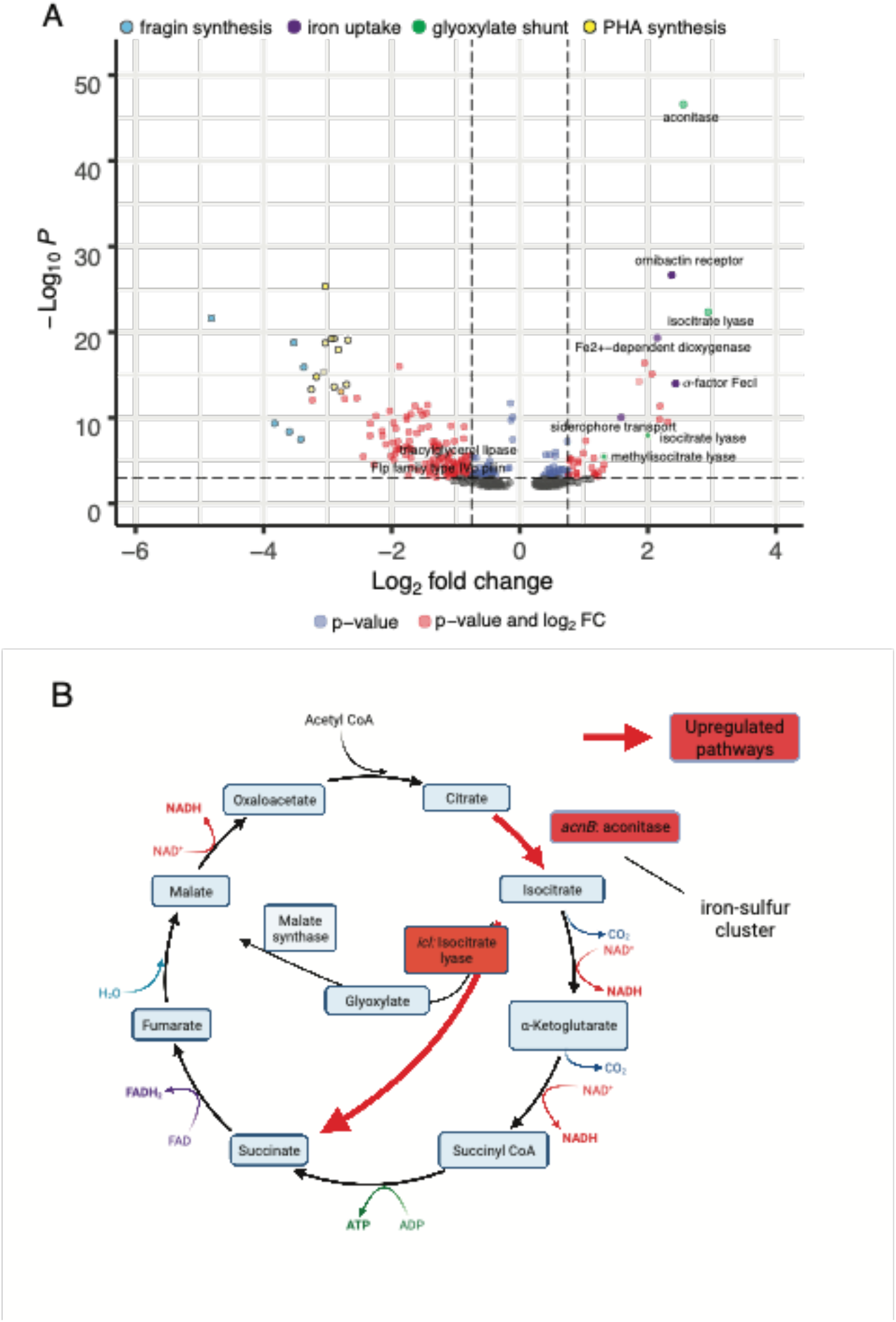
Genome-wide differences in expression between TilS N274Y and WT using RNAseq. A. Volcano plot of 431 genes with fold-change > 0.75 and/or p < 0.001 (denoted by dotted lines), with those meeting one or both criteria plotted in color. Most of the genes exhibiting the most significant changes in expression participate in four pathways, as shown. Comparisons between other mutants are shown in Supplementary Figure 5. B. Shared, upregulated enzymes in *tilS* and *tRNA*^*Ile2*^ mutants up-regulate the glyoxylate bypass, likely to preserve carbon (acetyl-CoA) for gluconeogenesis.

### *tilS* mutations are also selected in the Long-Term Evolution Experiment with *E. coli*

Our findings suggest that the enzymatic modification of tRNA^Ile2^ and/or a secondary function of TilS could experience selection in other evolution experiments with similar metabolic demands. The best studied and longest running evolution experiment is the Long-Term Evolution Experiment (LTEE), in which populations of *E. coli* have been propagated in minimal medium with glucose as sole carbon source for >70,000 generations. We hypothesized that this long-term selection for rapid growth and reduced lag phase on glucose could also select *tilS* mutants and searched published results of whole-genome sequencing of LTEE clones and populations for these mutations [36, 37]. We identified 13 unique mutations (12 nonsynonymous) in five independent populations (Supplementary Table 7); four of these became fixed (100% frequency) in three different populations (Figure 7). Based on the mutant activities shown here, we speculate that as many as three LTEE populations do not decode tRNA^Ile2^ as efficiently as the WT ancestral *E. coli*. This deficit could be significant because the *E. coli* genome encodes the AUA codon in the proteome in roughly 5 per 1000 codons, which is far greater than the rate of 0.6 per 1000 in

**Figure 7.**
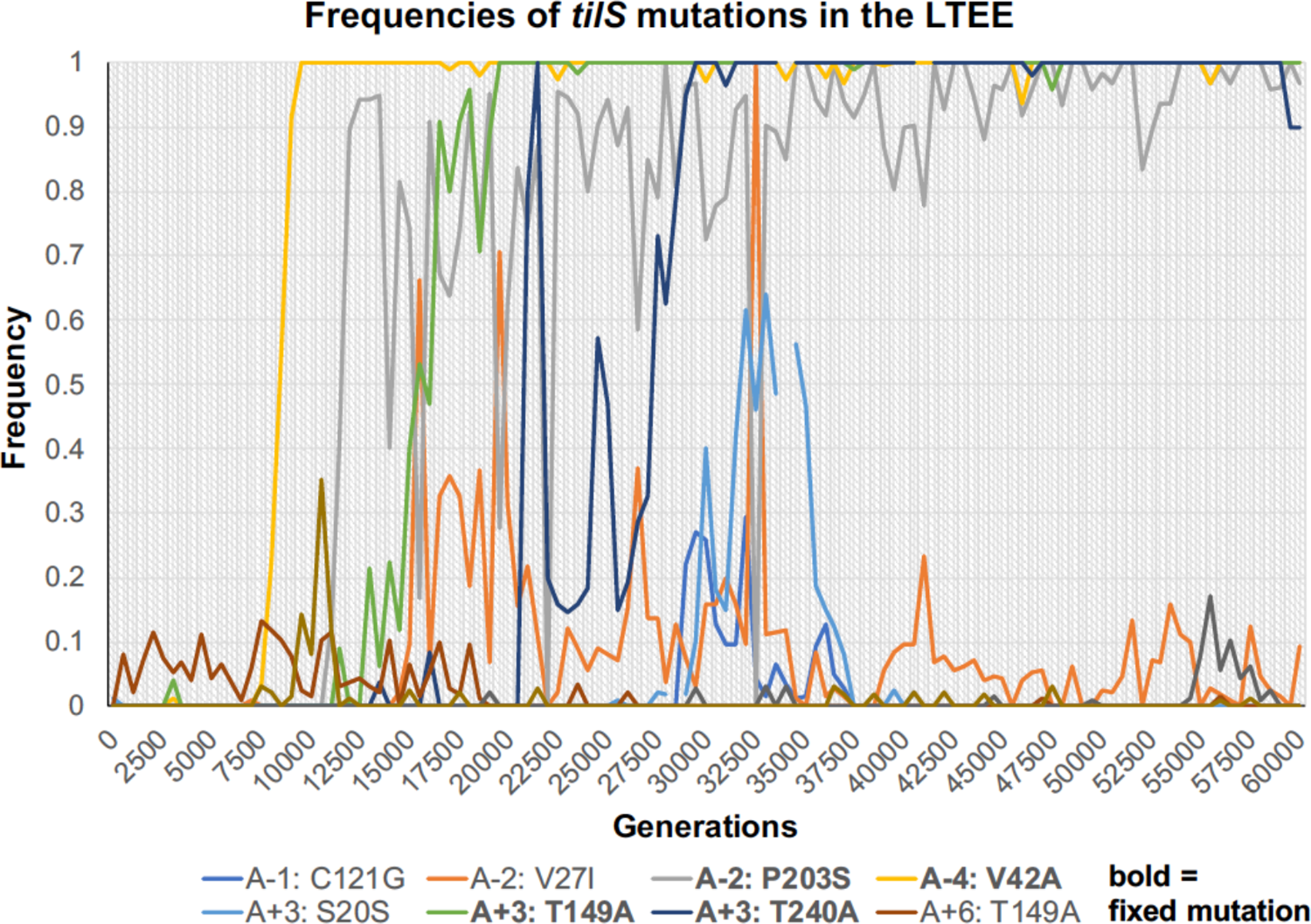
Frequencies of 10 *tilS* mutations in the LTEE that were detected at multiple time points during the experiment [36, 37]. Three other mutations were only detected in single clones (Supplementary Table 7). Four mutations denoted in bold reached 100% frequency: V42A in Ara-4, P203S in Ara-2, T149A in Ara+3, and T240A also in Ara+3.

*B. cenocepacia*. However, all fixed mutations arose in populations Ara-2, Ara-4, and Ara+3 that had evolved higher mutation rates, which increases the possibility that *tilS* mutations were not themselves beneficial but rather became fixed due to linkage with other more beneficial mutations [37]. On the other hand, identical T149A mutations arose independently in two populations, ultimately fixing in one of them (Supplementary Table 7). This precise molecular parallelism indicates they are adaptive, but the enzymatic or fitness phenotypes associated with these mutations remain to be characterized.

## Discussion

Research into the genetic causes of adaptation is advancing rapidly with the aid of innovative high-throughput screens and efficient whole-genome sequencing. Often, beneficial genotypes converge on a few pathways or genes that were unexpected from the experimental environment and its stressors [3, 29]. Selected mutants can reveal previously unknown genetic pathways to improved growth and disproportionately involve global regulators of expression [38, 39]. These possibilities motivated our evolution experiments that included a reporter of selective sweeps to capture the earliest adaptive mutants in a simple laboratory environment medium with galactose as the sole carbon source. In two experiments with different *B. cenocepacia* strains, we identified 32 genetically distinct mutants associated with increased fitness, and remarkably 11 of them carried mutations in the TilS pathway, either in *tilS* gene itself (n=7) or in its substrate *tRNA*^*Ile2*^ gene (n=4). All characterized *tilS* mutations reduced the known catalytic function of the TilS enzyme, the lysidinylation of the tRNA^Ile2^ anticodon that is essential for faithful and efficient translation of the AUA codon [18]. Because prior research demonstrated that a partial knockdown of lysidine formation drastically hinders AUA decoding *in vivo* [15], our discovery of >10-fold reduced lysidine production by evolved mutants suggests that their translation of AUA codons is significantly impaired. Yet these mutations improve fitness under our experimental conditions by reducing the lag phase prior to exponential growth (Figures 2,3), suggesting that at least under certain conditions, sacrificing translational fidelity and efficiency can be beneficial for growth. Single SNPs in *tilS* also dramatically influence the global transcriptome (Figure 6) and indicate that fitness gains are associated with increased usage of the glyoxylate bypass and iron scavenging and suppressed production of an antifungal secondary metabolite and PHA polymers. How mutated TilS or unmodified tRNA^Ile2^ alter expression of these pathways and enable earlier exponential growth (Figure 3) remains unclear and we discuss some possibilities below. However, the convergent evolution of mutations in both *tilS* and *tRNA*^*Ile2*^ suggests that interference with tRNA^Ile2^ lysidinylation was strongly favored.

Several studies have attempted to mutate or delete *tilS* from varied model organisms without success, except for the notable case of a *tilS* deletion mutant of *B. subtilis* that was suppressed by a co-occurring mutation in the anticodon of tRNA^Ile2^ from CAT to TAT [40]. This experimental evidence that *tilS* is an essential gene is supported by its nearly universal conservation in bacterial genomes, save for a few that encode tRNA^Ile2^ with a TAT anticodon [41]. However, the usage frequency of the AUA codon varies widely despite being relatively rare, from 0.6 AUA codons per 1000 in G/C-rich genomes like *B. cenocepacia* HI2424 to ∼27 AUA codons per 1000 in G/C-poor genomes like *Campylobacter* species [42]. It seems likely that varied demand for AUA decoding as a function of overall genome G/C content influences the essentiality of lysidinylation by TilS and could have facilitated the disruption of the TilS:tRNA^Ile2^ reaction in *B. cenocepacia*. Nonetheless, our discovery of 12 nonsynonymous mutations and 4 fixed mutations in the *E. coli* LTEE indicates that an organism with intermediate genome usage of AUA (5 per 1000) can still tolerate mutations in *tilS*. Further, the dynamics of the 10 LTEE mutations that persisted at detectable frequencies over thousands of generations combined with the fact that identical mutations arose in two independent lineages suggest that mutated TilS was also advantageous during long-term selection for rapid growth in minimal glucose medium (Figure 7). Together, the experimental selection of mutated *tilS* in two different systems demonstrates that partial loss-of-function mutations of this enzyme are not only tolerated, but even subject to positive selection under conditions of extreme metabolic flux that cause redox imbalance and demands for gluconeogenesis. Nevertheless, the fact that no deletion or premature stop mutations were selected in these experiments strongly indicates that translation of an intact TilS enzyme remains essential. The extreme pleiotropic effects of selected mutants on growth dynamics, resource usage, and global transcription point towards a yet-undiscovered moonlighting function for TilS, perhaps by binding other RNAs that regulate central metabolism and/or suppress production of secondary metabolites (Figure 6).

We considered several explanations for the fitness advantages of *tilS* and *tRNA*^*Ile2*^ mutations. First, we reasoned that genes enriched in rare AUA codons may have been selected for suppressed translation that an efficient TilS enzyme would otherwise enable. This form of attenuation by slowed translation speed has several precedents [43]. However, the *B. cenocepacia* genome has only 20 clearly predicted genes with 4 or more AUA codons, and only ∼1100 AUA codons in the entire 7.8 Mbp genome distributed across ∼800 genes [44]. Nonetheless, some of these 20 genes could play a role in modulating redox stress or metabolic regulation, such that their altered translation could influence fitness. Yet no mutations or transcriptional differences were observed in this gene set, implying that any resulting differences would need to be studied by comparing protein levels. Specifically, we predict that AUA codons would specify misincorporation of Met instead of Ile in mutant genotypes more than typical background translation error, which we are currently measuring.

Second, we considered that lysine levels became a limiting resource and their usage to modify tRNA^Ile2^ acted as a regulatory signal, however infrequent this reaction. The fact that supplementation with lysine or other amino acids that are readily converted to lysine complemented the WT fitness deficit supports this explanation (Figure 2). Further, mutant TilS enzymes exhibit reduced production of lysidine, and hence reduced consumption of lysine (Figure 5). One possible source of lysine demand is the need to buffer an acidic cytosol or periplasm caused by organic acid byproducts of growth using the Entner-Doudoroff pathway (Figure 3) [45]. In response to this acid stress, many microbes produce polyamines by decarboxylating amino acids like lysine and arginine into cadaverine, putrescine, or spermidine. These polyamines both directly buffer pH and interact with porins to prevent further proton uptake [46, 47], and research is mounting that they may have broader regulatory roles by activating the stringent response via the alarmone ppGpp [48]. We tested this model by quantifying levels of amino acids in WT and mutant strains grown in our selective conditions by HPLC, and we also attempted to measure cadaverine. However, no significant differences in amino acid or polyamine levels were observed repeatably, suggesting that any interactions with polyamine production, should they exist, are transient and/or limited to the lag phase of growth distinguishing WT and mutants, where cell density may have been too low for these measurements.

Third, it is possible that TilS, in its role as a tRNA-modifying enzyme, binds other RNAs relevant for emergence from lag phase under these experimental conditions. Several major differences in mutant transcriptomes were identified that are not simple reflections of earlier exponential growth, given that we harvested cells at similar density and growth rate. Two major pathways that were upregulated seem to be the best candidates for such interactions (Figure 6): activation of the glyoxylate bypass to preserve carbon backbones from the TCA cycle for biomass production, or increased capacity for iron uptake that is a hallmark for entering exponential growth [49]. Notably, the most upregulated gene in the TilS N274Y mutant is aconitase, which is an iron-sulfur cluster enzyme that leads to the glyoxylate bypass as well as TCA. This change, combined with increased siderophore activity, suggests that mutant TilS may enable acquisition of trace metals to exit lag sooner. Future work will examine this potential secondary function of TilS. Fourth, it is possible that the modification state of tRNA^Ile2^ may act as a metabolic sensor in an unknown pathway. It is becoming appreciated that tRNAs can govern metabolism by linking their decoding roles with biosynthetic processes [43]. Notable examples include the *miaB* gene that thiomethylates adenosine residues in tRNAs with NNA anticodons and also influences sporulation and antibiotic production in *Streptomyces* [50]. In addition, the *miaE* gene that produces ms^2^i^6^A nucleoside modifications also influences aerobic growth on dicarboxylic acids in *Salmonella* [51]. The most promising precedent may be a mutant of *tilS* in *Helicobacter pylori* that was associated with increased colonization of the mouse gut, an acidic environment, but this association was not explored further [52]. In summary, based on the findings presented here, either the unmodified tRNA^Ile2^ or the unoccupied TilS enzyme likley play roles in regulating central metabolism that extend beyond their role in translational accuracy, but the identity of these genetic partners still elude us.

To conclude, a screen for beneficial genetic mutants of *B. cenocepacia* growing in a simple, inadequately buffered growth medium containing galactose as sole carbon source repeatedly selected mutants that should in theory fail to properly decode the rare AUA codon. These mutants improved growth by reducing the duration of lag phase, resulting in a large competitive advantage, and their advantages were specific to rapid, redox-imbalanced growth in medium lacking certain amino acids. TilS mutants also evolved in a long-running evolution experiment with *E. coli* in which exit from lag phase is also at a premium [53] and exhibited dynamics consistent with an adaptive role, together suggesting this tRNA modification pathway may influence fitness in diverse bacteria. The breadth of phenotypes altered by these mutations add to the growing evidence that the many modifications of tRNAs, each of which link to distinct pathways and essential building blocks, enable exquisite sensing of the cellular metabolic state.

## Acknowledgments

We thank Rachel Staples and David Morejon for isolating and beginning to characterize these mutants 10 years ago, Christopher Marshall and Nara Lee for helpful discussion, and Daniel Snyder for technical assistance. This research was supported by NSF MCB-1818131 to RWA and VSC.

---

**Supplementary Figure 1.**
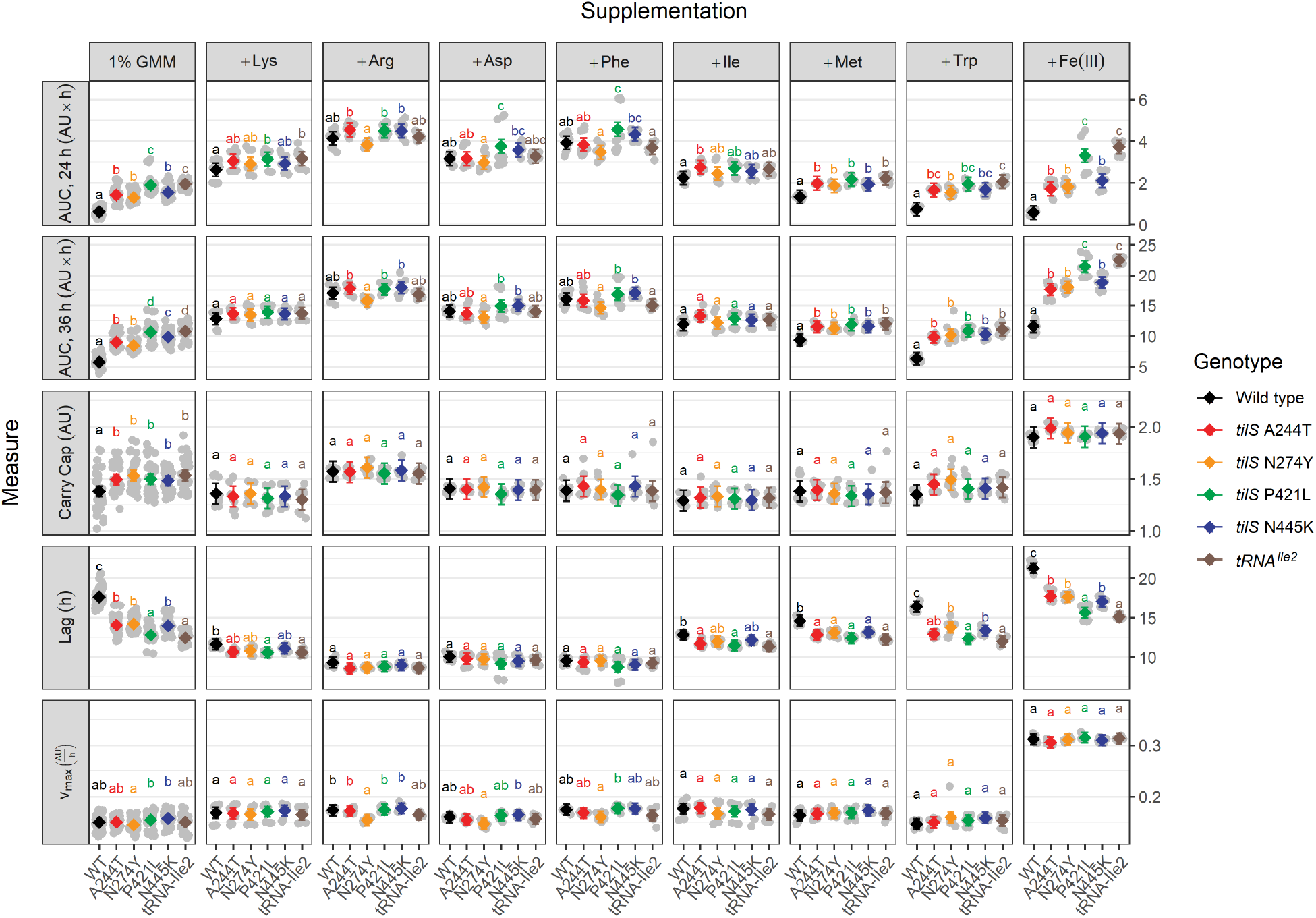
Effects of supplements to GMM (column 1) on components of fitness that differentiate tilS mutants from WT. AUC = area under the curve, for 24h or 36. Lag, carrying capacity (carry cap) and maximum growth rate (v_max_) are inferred from the growth curve (see methods). Means and confidence intervals are in blue (WT) and black (mutants), with individual observations in grey. Different letters group together related patterns of statistically significant differences, determined via post-hoc Šidák-corrected pairwise means testing with a cut-off of 0.05 following a two-way ANOVA. Quantitative differences in fitness measures (WT-mutant) are shown in boxes, and significant differences are shaded by magnitude of difference. AU = absorbance units at OD600. h = hours. Lys = lysine. Asp = aspartate. Met = Methionine.

**Supplementary Figure 2.**
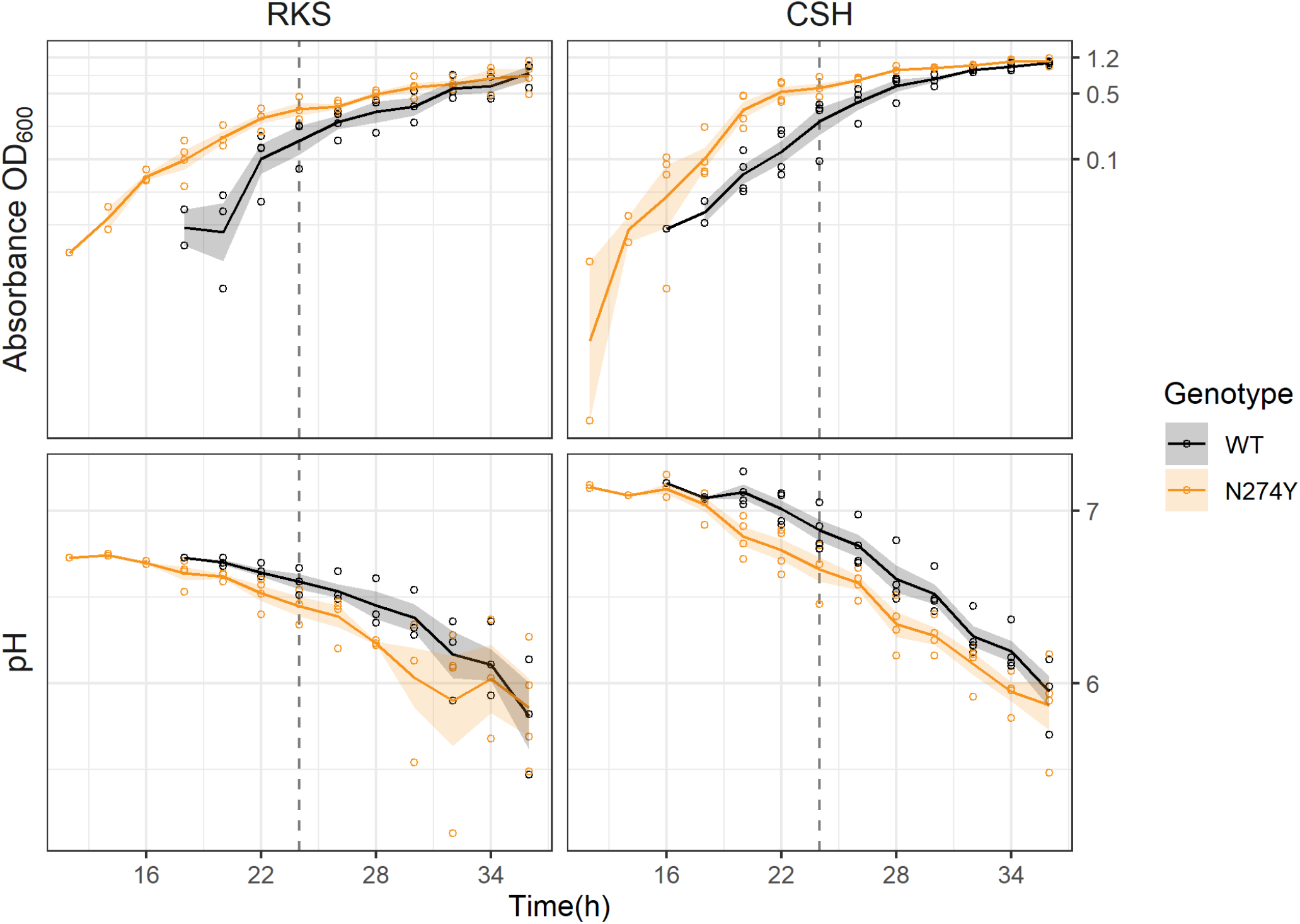
Acidification of media with *B. cenocepacia* growth. Representative *tilS* mutant N274Y (orange) reaches higher OD than WT (black) *B. cenocepacia* by the 24H transfer period (dotted line) in selective RKS medium and in better buffered CSH medium, while acidifying the medium significantly more consequently. Lines and ribbons are means and standard error by time point.

**Supplementary Figure 3.**
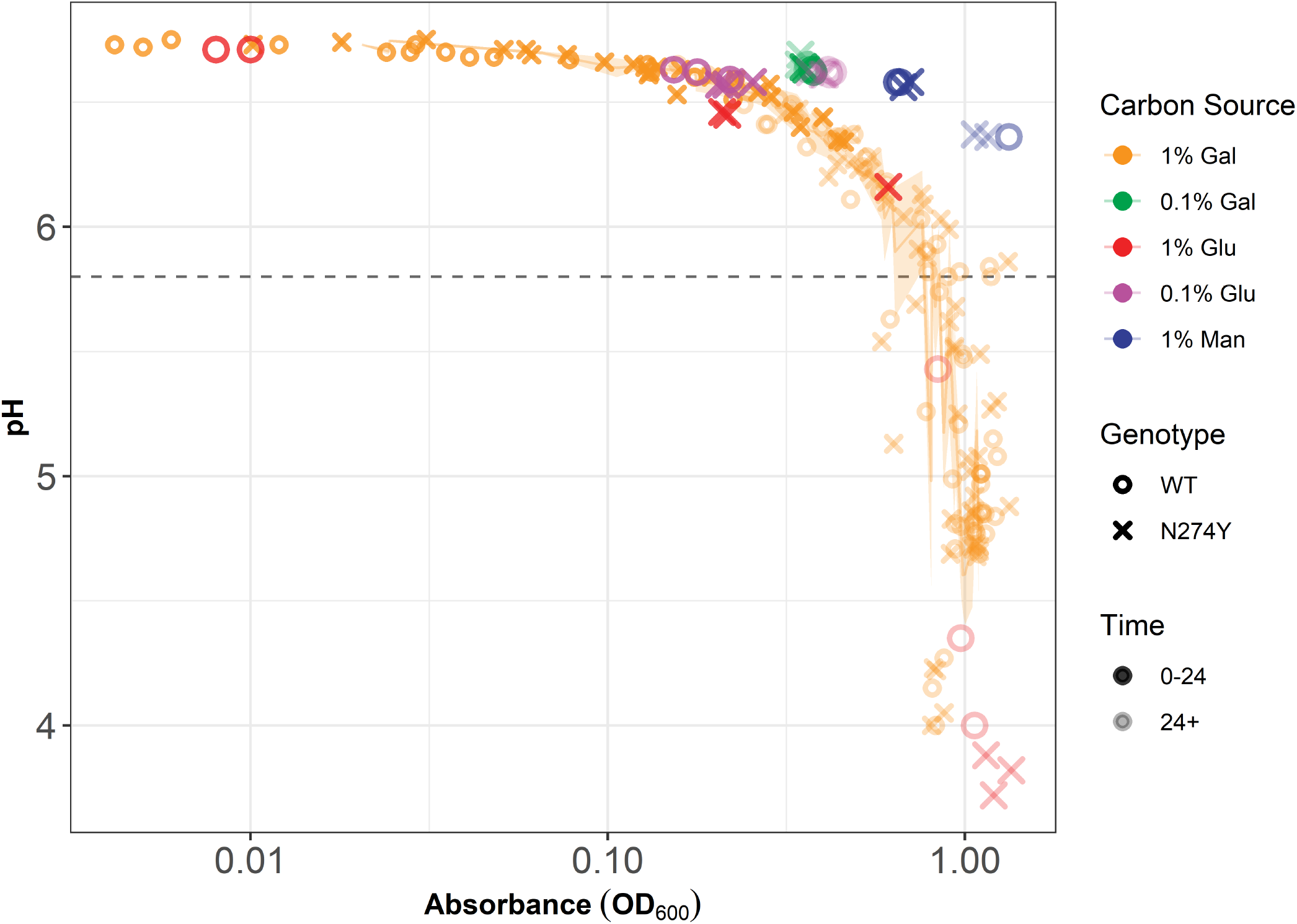
Relationship between growth, as measured by absorbance, and pH, in different media. Overflow metabolism in 1% galactose or 1% glucose leads to progressive acidification of the medium over time by both WT and representative *tilS* mutant N274Y, which follow the same relationship between these variables. Growth in 1.0% mannose, 0.1% galactose, or 0.1% glucose do not acidify the medium at the same rate as 1.0% galactose. Dotted line = predicted buffering capacity of the medium.

**Supplementary Figure 4.**
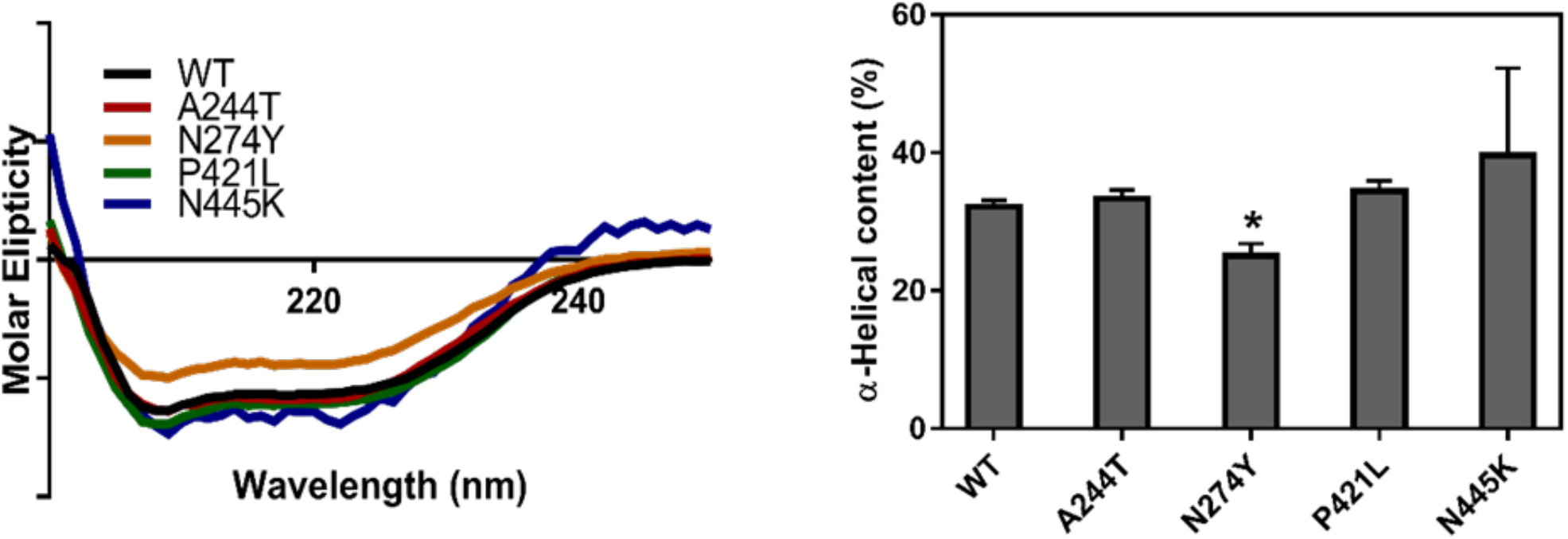
Circular dichroism of TilS variants. Spectra were collected in 20 mM Tris-HCl (pH 7.8) buffer. A. Spectra represent the averages of experimental triplicates each of which were analyzed from technical triplicates. B. Bar graph shows the determined α-helical content of each variant; error bars indicate the standard deviation from three experimental replicates and * indicates statistically different composition (P<0.05) compared to wild-type TilS using an ANOVA statistical comparison.

**Supplementary Figure 5.**
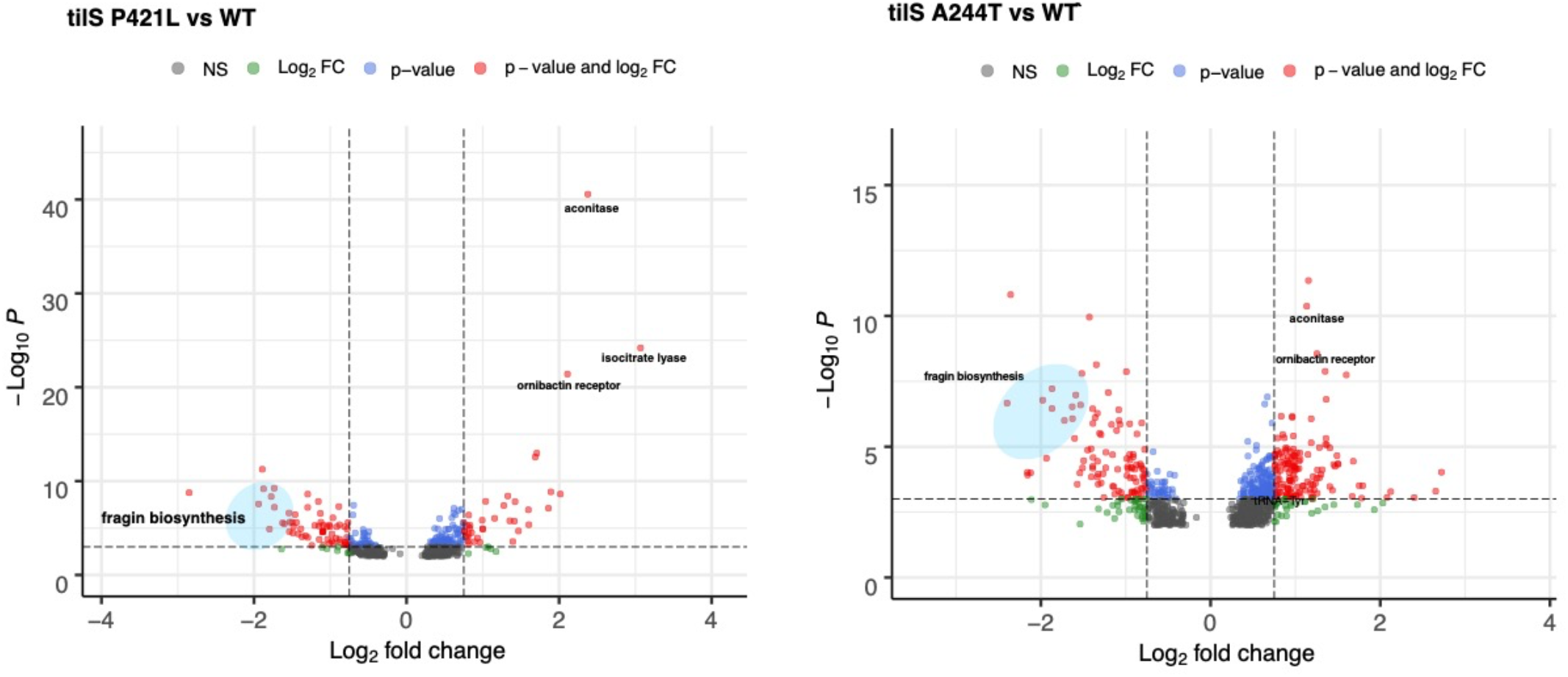
Volcano plots of comparisons between the transcriptome of WT and *tilS* mutants A244T and P421L. Key similarities in strongly upregulated and downregulated genes with Figure 6 (N274Y vs WT) are highlighted. Tables of these and other primary analyses are available at http://github.com/vscooper/tilS

**Supplementary Table 1.**
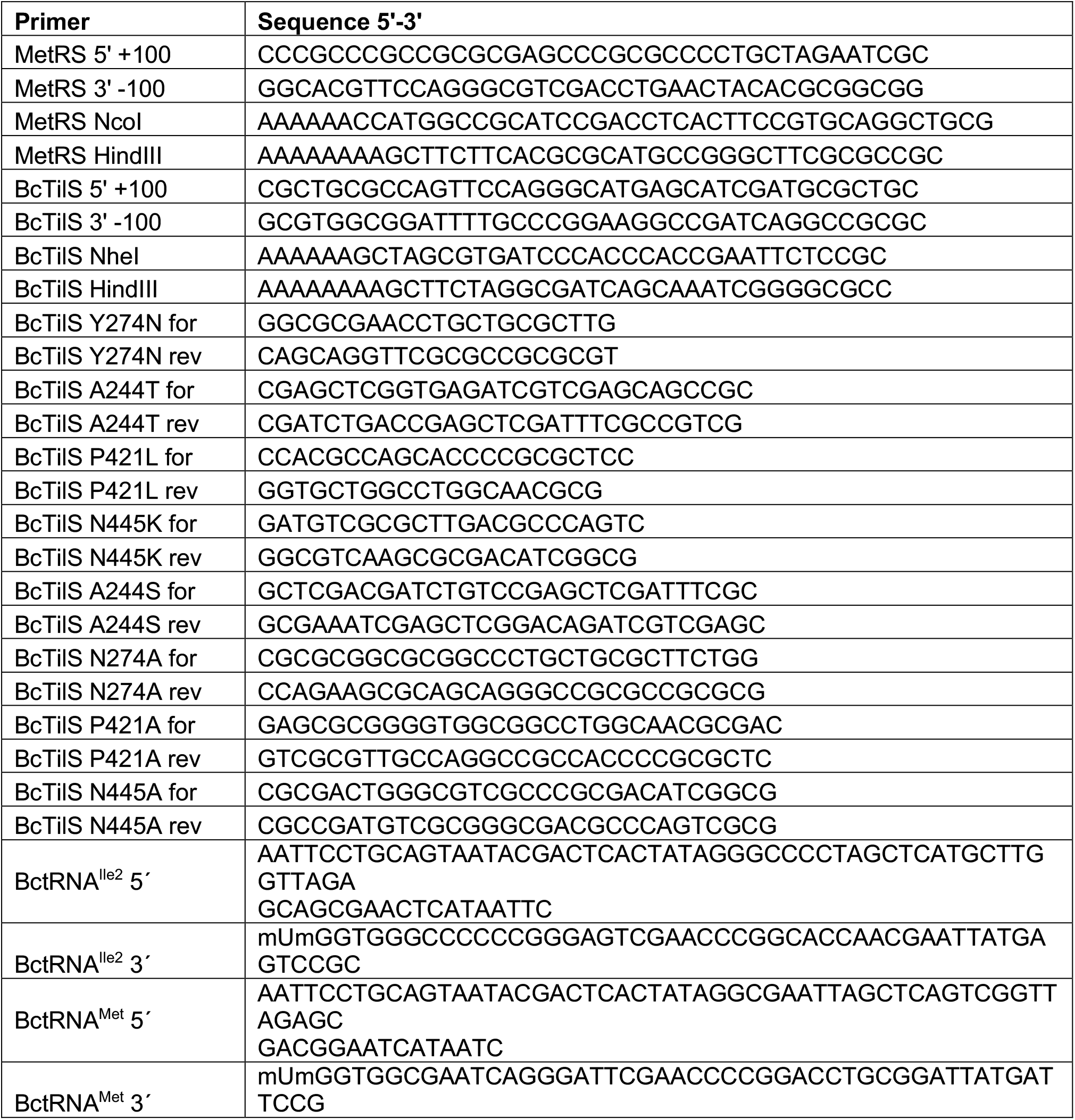
Primers used for cloning and mutagenesis.

**Supplementary Table 2.** Trace elements to mimic the water used for the original evolution experiment in which mutants were selected.

| Element | 1X Conc.<br>($\mu\text{g/L}$) | Source<br>Compound | |
| --- | --- | --- | --- |
| Calcium | 40 | $\text{CaCl}_2$ | |
| Manganese | 0.3 | $\text{MnSO}_4$ | |
| Potassium | 11 | KBr |  |
| Sodium | 25 | $\text{Na}_2\text{SiO}_3$ | |
| Zinc | 50 | $\text{ZnSO}_4$ | |
| Cobalt | 0.044 | $\text{CoCl}_2 \cdot (\text{H}_2\text{O})_6$ | |
| Copper | 0.58 | $\text{CuSO}_4$ | |

**Supplementary Table 3.** Relative fitness of mutants captured during experimental evolution using marker deflection, assessed over 24h vs WT HI2424.

| Mutant | Relative Fitness $r$ | 95% ci |
| --- | --- | --- |
| 1 | 0.629 | 0.315 |
| 2 | 1.043 | 0.230 |
| 3 | 1.399 | 0.436 |
| 4 | 1.102 | 0.239 |
| 5 | 1.065 | 0.611 |
| 6 | 0.843 | 0.158 |
| 7 | 2.037 | 0.313 |
| 8 | 1.299 | 0.157 |
| 9 | 1.380 | 0.128 |
| 10 | 1.433 | 0.775 |
| 12 | 1.512 | 0.435 |
| 13 | 0.654 | 0.551 |
| 14 | 1.740 | 0.293 |
| 15 | 0.877 | 0.287 |
| 16 | 0.499 | 0.687 |
| 17 | 0.460 | 0.657 |
| 19 | 0.882 | 0.450 |

**Supplementary Table 4.** All mutations identified by WGS in 17 *B. cenocepacia* mutants selected for improved fitness in minimal medium containing galactose as sole carbon source. Clones 11 and 18 had no identifiable mutations and were excluded. Mutants in bold are the focus of this study.

| Mutant | Locus | Annotation | Position | Amino Acid |
| --- | --- | --- | --- | --- |
| **1** | **Bcen2424_2075**<br>**(*tilS*)** | **tRNA(Ile)-lysidine synthetase** | **2305668** | **R208C (CGC→TGC)** |
| 1 | Bcen2424_0661<br>(*eda*) | KDPG_aldolase | 736233 | P207L (CCC→CTC) |
| 1 | Bcen2424_3218 | short-chain dehydrogenase/reductase SDR | 59305 | Q35H (CAG→CAT) |
| 1 | Bcen2424_4619 | RND efflux system outer membrane lipoprotein | 1597143 | P267L (CCG→CTG) |
| 2 | Bcen2424_2426<br>(*ppc*) | phosphoenolpyruvate carboxylase | 2691740 | T41T (ACG→ACC) |
| 2 | Bcen2424_6531 | acriflavine resistance protein | 756381 | V225A (GTC→GCC) |
| 3 | Bcen2424_2426 | phosphoenolpyruvate carboxylase | 2,691,756 | D47N (GAC→AAC) |
| 3 | Bcen2424_5220 | ATPase central domain-containing protein | 2,277,234 | L202P (CTC→CCC) |
| 4 | *hemC* | porphobilinogen deaminase | 2,691,597 | A7G (GCT→GGT) |
| **5** | ***tilS*** | **tRNA(Ile)-lysidine synthetase** | **2,304,955** | **N445K (AAT→AAG)** |
| **6** | ***tilS*** | **tRNA(Ile)-lysidine synthetase** | **2,305,028** | **P421L (CCG→CTG)** |
| 7 | Bcen2424_6711 | major facilitator transporter | 948,195 | Δ3 bp (1110-1112/1329) |
| **8** | ***tilS*** | **tRNA(Ile)-lysidine synthetase** | **2,305,470** | **N274Y (AAC→TAC)** |
| 9 | Bcen2424_2627<br>(*pyk*) | pyruvate kinase | 2911524 | Δ1 bp (486/1437) |
| 9 | Bcen2424_0478 | hypothetical protein | 533982 | D199E (GAC→GAG) |
| 9 | Bcen2424_4159 | major facilitator transporter | 1073395 | L14P (CTC→CCC) |
| 10 | Bcen2424_2426 | *ppc*: phosphoenolpyruvate carboxylase | 2,691,740 | T41T (ACG→ACT) |
| 10 | Bcen2424_0378 /<br>*engB* | delta-aminolevulinic acid dehydratase/ribosome biogenesis GTP-binding protein YsxC | 423,603 | intergenic (-180/+73) |
| 12 | Bcen2424_0610 | UDP-N-acetylmuramate | 680,525 | Q41* (CAG→TAG) |
| 12 | Bcen2424_4159 | major facilitator transporter | 1,073,395 | L14P (CTC→CCC) |
| **13** | ***tilS*** | **tRNA(Ile)-lysidine synthetase** | **2,305,560** | **A244T (GCC→ACC)** |
| 14 | *ppc* | phosphoenolpyruvate carboxylase | 2,691,740 | T41T (ACG→ACT) |
| **15** | **Bcen2424_R0075** | **tRNA-Ile<sup>2</sup>** | **753458** | **noncoding: A79G** |
| 16 | Bcen2424_R0023 | tRNA-Lys | 900,484 | noncoding:<br>(TGGGAGGG)1→2<br>(62/77 nt) |
| 17 | Bcen2424_0524 /<br>Bcen2424_0525 | GTP-dependent nucleic acid-binding protein EngD/ubiquinone biosynthesis hydroxylase family protein | 584933 | intergenic (-119/-130) |
| 17 | Bcen2424_3118 | organic radical activating protein-like protein | 3433117 | I113V (ATC→GTC) |
| 17 | Bcen2424_5939 | response regulator receiver modulated metal dependent phosphohydrolase | 73960 | A464A (GCA→GCC) |
| **19** | ***tilS*** | **tRNA(Ile)-lysidine synthetase** | **2,305,560** | **A244T (GCC→ACC)** |

**Supplementary Table 5.** All mutations identified by WGS in 15 adaptive mutants of the *B. cenocepacia* “Step 7” clone that had been preadapted in GMM under biofilm-selective conditions. Mutants in bold are the focus of this study. Clones 1 and 15 had no identifiable mutations and were excluded.

| Mutant | Locus | Annotation | Position | Amino Acid |
| --- | --- | --- | --- | --- |
| **2** | **Bcen2424_R0075** | **tRNA-Ile<sup>2</sup>** | **753394** | **noncoding: 15/79 nt** |
| 3 | *hemC* | porphobilinogen deaminase | 2691276 | P114Q (G->T) |
| 4 | *ppc* | phosphoenolpyruvate carboxylase | 2691756 | D47Y (G>T) |
| 5 | *ppc* | phosphoenolpyruvate carboxylase | 2691756 | D47Y (G>T) |
| 5 | Bcen2424_2427 / Bcen2424_2428 | type 11 methyltransferase / XRE family transcriptional regulator | 2695639 | -105/-122 (C>G) |
| 6 | Bcen2424_0547 | hypothetical | 616106 | Q28R (A>G) |
| 6 | Bcen2424_3441 | hypothetical | 271700 | V178G (T>G) |
| 6 | *pyk* | pyruvate kinase | 2911758 | 252/1437 (Δ1 bp) |
| 7 | Bcen2424_4077 | enoyl-CoA hydratase | 972475 | A189G (C>G) |
| 7 | *ppc* | phosphoenolpyruvate carboxylase | 2691740 | T41T (G>C) |
| **8** | **Bcen2424_R0075** | **tRNA-Ile<sup>2</sup>** | **753442** | **noncoding: 63/79 nt (T>G)** |
| 9 | Bcen2424_5460 / Bcen2424_5461 | Hsp90 ATPase 1 | 2559014 | +108/-12 (G>A) |
| 9 | *ppc* | phosphoenolpyruvate carboxylase | 2691630 | G5R (G>A) |
| 10 | Bcen2424_4062 | L-asparaginase II | 956271 | A202G (C>G) |
| 10 | Bcen2424_1349 | periplasmic binding | 1488451 | D214E (C>A) |
| **10** | **Bcen2424_R0075** | **tRNA-Ile<sup>2</sup>** | **753442** | **Non-coding 63/79 nt (T>G)** |
| 11 | Bcen2424_4498 | hypothetical protein | 1450756 | K455K (C>T) |
| 11 | Bcen2424_5069 / Bcen2424_5070 | major facilitator transporter / porin | 2093505 | -145/+98 (C>G) |
| 11 | Bcen2424_0785 | N-acetyltransferase | 891452 | A106P (G>C) |
| **11** | **Bcen2424_R0075** | **tRNA-Ile<sup>2</sup>** | **753442** | **Non-coding 63/79 nt (T>G)** |
| 12 | *ppc* | phosphoenolpyruvate carboxylase | 2691630 | G5R (G>C) |
| 13 | Bcen2424_5091 | HlyD family protein | 2121571 | V325V (C>G) |
| 13 | *ppc* | phosphoenolpyruvate carboxylase | 2691631 | G5E (G>A) |
| 13 | Bcen2424_3038 | urea amidolyase | 3351696 | A246P (G>C) |
| 14 | *ppc* | phosphoenolpyruvate carboxylase | 2691625 | S3C (C>G) |
| 16 | Bcen2424_2914 | hypothetical protein | 3218565 | K93N (A>C) |
| 16 | *ppc* | phosphoenolpyruvate carboxylase | 2691630 | G5R (G>A) |
| 17 | Bcen2424_6582 | MgtC/SapB | 806881 | A114A (T>C) |
| 17 | Bcen2424_2426 | phosphoenolpyruvate carboxylase | 2691757 | D47V (A>T) |
| 17 | Bcen2424_3038 | urea amidolyase | 3351696 | A246P (G>C) |

**Supplementary Table 6.** Error represents the standard error of the mean for each parameter as determined from three replicates. NM, not measurable.

|  | WT | A244T | N274Y | P421L | N445K |
| --- | --- | --- | --- | --- | --- |
| Lysidinylation rate (nM/min) | $5.8 \pm 0.8$ | $0.17 \pm 0.01$ | $0.08 \pm 0.01$ | NM | $0.9 \pm 0.1$ |
| Catalytic loss (fold) | 1 | 34 | 72 | >100 | 6 |
| tRNA<sup>Ile2</sup> K<sub>d</sub> (μM) | $2.6 \pm 0.3$ | $4.6 \pm 0.7$ | $3.3 \pm 0.4$ | $4.5 \pm 0.5$ | $3.1 \pm 0.3$ |
| Relative cellular lysidinylation | 1 | $0.3 \pm 0.09$ | $0.3 \pm 0.1$ | $0.1 \pm 0.05$ | $0.3 \pm 0.02$ |

**Supplementary Table 7.** *tilS* mutations identified in the Long-Term Evolution Experiment with *E. coli* ([36, 37]

| Population | Gene position | Allele | Annotation | Maximum observed frequency |
| --- | --- | --- | --- | --- |
| Ara-1 | 361 | T->G | C121G | 0.271 |
| Ara-2 | 79 | G->A | V27I | 0.71 |
| Ara-2 | 607 | C->T | P203S | 1 |
| Ara-4 | 125 | T->C | V42A | 1 |
| Ara-4 | 949 | A->G | S317G | 1 clone |
| Ara-4 | 101 | A->G | Q34R | 1 clone |
| Ara+3 | 60 | C->T | S20S | 0.61 |
| Ara+3 | 220 | C->A | P74T | 1 clone |
| Ara+3 | 445 | A->G | T149A | 1 |
| Ara+3 | 718 | A->G | T240A | 1 |
| Ara+6 | 445 | A->G | T149A | 0.132 |
| Ara+6 | 464 | T->G | L155W | 0.171 |
| Ara+6 | 1106 | T->G | V369G | 0.352 |

## References

1. Lang GI, Desai MM. The spectrum of adaptive mutations in experimental evolution. Genomics 2014; 104: 412–416.

2. Cooper VS. Experimental Evolution as a High-Throughput Screen for Genetic Adaptations. mSphere 2018; 3.

3. Bailey SF, Bataillon T. Can the experimental evolution programme help us elucidate the genetic basis of adaptation in nature? Molecular Ecology 2016; 25: 203–218.

4. Ostrowski EA, Woods RJ, Lenski RE. The genetic basis of parallel and divergent phenotypic responses in evolving populations of Escherichia coli. Proceedings Biological sciences / The Royal Society 2008; 275: 277–284.

5. Turner CB, Marshall CW, Cooper VS. Parallel genetic adaptation across environments differing in mode of growth or resource availability. Evolution Letters 2018; 2: 355–367.

6. Bolnick DI, Barrett RDH, Oke KB, Rennison DJ, Stuart YE. (Non)Parallel Evolution. Annual Review of Ecology, Evolution, and Systematics 2018; 49: null.

7. Wiser MJ, Ribeck N, Lenski RE. Long-Term Dynamics of Adaptation in Asexual Populations. Science 2013; 342: 1364–1367.

8. Cooper VS, Schneider D, Blot M, Lenski RE. Mechanisms causing rapid and parallel losses of ribose catabolism in evolving populations of Escherichia coli B. J Bacteriol 2001; 183: 2834–2841.

9. Cooper TF, Rozen DE, Lenski RE. Parallel changes in gene expression after 20,000 generations of evolution in Escherichia coli. Proc Natl Acad Sci U S A 2003; 100: 1072–7.

10. Consuegra J, Plucain J, Gaffé J, Hindré T, Schneider D. Genetic Basis of Exploiting Ecological Opportunity During the Long-Term Diversification of a Bacterial Population. J Mol Evol 2017; 85: 26–36.

11. Rozen DE, de Visser J, Gerrish PJ. Fitness effects of fixed beneficial mutations in microbial populations. Curr Biol 2002; 12: 1040–1045.

12. Hegreness M, Kishony R. Analysis of genetic systems using experimental evolution and whole-genome sequencing. Genome Biology 2007; 8: 201.

13. Cooper VS, Staples RK, Traverse CC, Ellis CN. Parallel evolution of small colony variants in Burkholderia cenocepacia biofilms. Genomics 2014; 104: 447–52.

14. Dillon MM, Rouillard NP, Van Dam B, Gallet R, Cooper VS. Diverse phenotypic and genetic responses to short-term selection in evolving Escherichia coli populations. Evolution 2016; 70: 586–599.

15. Soma A, Ikeuchi Y, Kanemasa S, Kobayashi K, Ogasawara N, Ote T, et al. An RNA-Modifying Enzyme that Governs Both the Codon and Amino Acid Specificities of Isoleucine tRNA. Molecular Cell 2003; 12: 689–698.

16. Gerdes SY, Scholle MD, Campbell JW, Balázsi G, Ravasz E, Daugherty MD, et al. Experimental determination and system level analysis of essential genes in Escherichia coli MG1655. J Bacteriol 2003; 185: 5673–5684.

17. Baba T, Ara T, Hasegawa M, Takai Y, Okumura Y, Baba M, et al. Construction of Escherichia coli K-12 in-frame, single-gene knockout mutants: the Keio collection. Mol Syst Biol 2006; 2.

18. Suzuki T, Miyauchi K. Discovery and characterization of tRNAIle lysidine synthetase (TilS). FEBS Lett 2010; 584: 272–277.

19. Muramatsu T, Nishikawa K, Nemoto F, Kuchino Y, Nishimura S, Miyazawa T, et al. Codon and amino-acid specificities of a transfer RNA are both converted by a single post-transcriptional modification. Nature 1988; 336: 179–181.

20. LiPuma JJ, Spilker T, Coenye T, Gonzalez CF. An epidemic Burkholderia cepacia complex strain identified in soil. The Lancet 2002; 359: 2002–2003.

21. Cooper VS, Staples RK, Traverse CC, Ellis CN. Parallel evolution of small colony variants in Burkholderia cenocepacia biofilms. Genomics 2014; 104: 447–452.

22. Chevin L-M. On measuring selection in experimental evolution. Biology Letters 2011; 7: 210–213.

23. M9 minimal medium (standard). Cold Spring Harb Protoc 2010; 2010: pdb.rec12295.

24. Nakanishi K, Bonnefond L, Kimura S, Suzuki T, Ishitani R, Nureki O. Structural basis for translational fidelity ensured by transfer RNA lysidine synthetase. Nature 2009; 461: 1144–8.

25. Nakanishi K, Fukai S, Ikeuchi Y, Soma A, Sekine Y, Suzuki T, et al. Structural basis for lysidine formation by ATP pyrophosphatase accompanied by a lysine-specific loop and a tRNA-recognition domain. Proceedings of the National Academy of Sciences 2005; 102: 7487–7492.

26. Edwards AM, Addo MA, Dos Santos PC. tRNA Modifications as a Readout of S and Fe-S Metabolism. In: Dos Santos PC (ed). Fe-S Proteins: Methods and Protocols. 2021. Springer US, New York, NY, pp 137–154.

27. Mhatre E, Snyder DJ, Sileo E, Turner CB, Buskirk SW, Fernandez NL, et al. One gene, multiple ecological strategies: A biofilm regulator is a capacitor for sustainable diversity. PNAS 2020; 117: 21647–21657.

28. Ashkenazy H, Abadi S, Martz E, Chay O, Mayrose I, Pupko T, et al. ConSurf 2016: an improved methodology to estimate and visualize evolutionary conservation in macromolecules. Nucleic Acids Research 2016; 44: W344–W350.

29. Traverse CC, Mayo-Smith LM, Poltak SR, Cooper VS. Tangled bank of experimentally evolved Burkholderia biofilms reflects selection during chronic infections. PNAS 2013; 110: E250–E259.

30. Silva IN, Ramires MJ, Azevedo LA, Guerreiro AR, Tavares AC, Becker JD, et al. Regulator LdhR and d-Lactate Dehydrogenase LdhA of Burkholderia multivorans Play Roles in Carbon Overflow and in Planktonic Cellular Aggregate Formation. Appl Environ Microbiol 2017; 83: e01343–17.

31. Turner CB, Buskirk SW, Harris KB, Cooper VS. Negative frequency-dependent selection maintains coexisting genotypes during fluctuating selection. Mol Ecol 2020; 29: 138–148.

32. Kobayashi K, Ehrlich SD, Albertini A, Amati G, Andersen KK, Arnaud M, et al. Essential Bacillus subtilis genes. Proc Natl Acad Sci U S A 2003; 100: 4678–4683.

33. Rolfe MD, Rice CJ, Lucchini S, Pin C, Thompson A, Cameron ADS, et al. Lag Phase Is a Distinct Growth Phase That Prepares Bacteria for Exponential Growth and Involves Transient Metal Accumulation. J Bacteriol 2012; 194: 686–701.

34. Dolan SK, Welch M. The Glyoxylate Shunt, 60 Years On. Annual Review of Microbiology 2018; 72: 309–330.

35. Jenul C, Sieber S, Daeppen C, Mathew A, Lardi M, Pessi G, et al. Biosynthesis of fragin is controlled by a novel quorum sensing signal. Nat Commun 2018; 9: 1297.

36. Good BH, McDonald MJ, Barrick JE, Lenski RE, Desai MM. The dynamics of molecular evolution over 60,000 generations. Nature 2017; 551: 45–50.

37. Barrick JE, Yu DS, Yoon SH, Jeong H, Oh TK, Schneider D, et al. Genome evolution and adaptation in a long-term experiment with Escherichia coli. Nature 2009; 461: 1243–7.

38. Maddamsetti R, Hatcher PJ, Green AG, Williams BL, Marks DS, Lenski RE. Core Genes Evolve Rapidly in the Long-Term Evolution Experiment with Escherichia coli. Genome Biology and Evolution 2017; 9: 1072–1083.

39. Venkataram S, Dunn B, Li Y, Agarwala A, Chang J, Ebel ER, et al. Development of a Comprehensive Genotype-to-Fitness Map of Adaptation-Driving Mutations in Yeast. Cell 2016; 166: 1585-1596.e22.

40. Fabret C, Dervyn E, Dalmais B, Guillot A, Marck C, Grosjean H, et al. Life without the essential bacterial tRNAIle2–lysidine synthetase TilS: a case of tRNA gene recruitment in Bacillus subtilis. Molecular Microbiology 2011; 80: 1062–1074.

41. Nilsson EM, Alexander RW. Bacterial wobble modifications of NNA-decoding tRNAs. IUBMB Life 2019; 71: 1158–1166.

42. Wan XF, Xu D, Kleinhofs A, Zhou J. Quantitative relationship between synonymous codon usage bias and GC composition across unicellular genomes. BMC Evol Biol 2004; 4: 19.

43. de Crécy-Lagard V, Jaroch M. Functions of Bacterial tRNA Modifications: From Ubiquity to Diversity. Trends in Microbiology 2021; 29: 41–53.

44. Nakamura Y, Gojobori T, Ikemura T. Codon usage tabulated from international DNA sequence databases: status for the year 2000. Nucleic Acids Res 2000; 28: 292.

45. Silva IN, Ramires MJ, Azevedo LA, Guerreiro AR, Tavares AC, Becker JD, et al. Regulator LdhR and d-Lactate Dehydrogenase LdhA of Burkholderia multivorans Play Roles in Carbon Overflow and in Planktonic Cellular Aggregate Formation. Appl Environ Microbiol 2017; 83.

46. Ma W, Chen K, Li Y, Hao N, Wang X, Ouyang P. Advances in Cadaverine Bacterial Production and Its Applications. Engineering 2017; 3: 308–317.

47. Samartzidou H, Mehrazin M, Xu Z, Benedik MJ, Delcour AH. Cadaverine Inhibition of Porin Plays a Role in Cell Survival at Acidic pH. J Bacteriol 2003; 185: 13–19.

48. Kanjee U, Gutsche I, Alexopoulos E, Zhao B, El Bakkouri M, Thibault G, et al. Linkage between the bacterial acid stress and stringent responses: the structure of the inducible lysine decarboxylase. The EMBO Journal 2011; 30: 931–944.

49. Bertrand RL. Lag Phase Is a Dynamic, Organized, Adaptive, and Evolvable Period That Prepares Bacteria for Cell Division. Journal of Bacteriology 2019; 201.

50. Sehin Y, Koshla O, Dacyuk Y, Zhao R, Ross R, Myronovskyi M, et al. Gene ssfg_01967 (miaB) for tRNA modification influences morphogenesis and moenomycin biosynthesis in Streptomyces ghanaensis ATCC14672. Microbiology (Reading) 2019; 165: 233–245.

51. Mathevon C, Pierrel F, Oddou J-L, Garcia-Serres R, Blondin G, Latour J-M, et al. tRNA-modifying MiaE protein from Salmonella typhimurium is a nonheme diiron monooxygenase. Proceedings of the National Academy of Sciences 2007; 104: 13295–13300.

52. Castillo AR, Woodruff AJ, Connolly LE, Sause WE, Ottemann KM. Recombination-Based In Vivo Expression Technology Identifies Helicobacter pylori Genes Important for Host Colonization. Infection and Immunity 2008.

53. Lenski RE, Mongold JA, Sniegowski PD, Travisano M, Vasi F, Gerrish PJ, et al. Evolution of competitive fitness in experimental populations of E. coli: what makes one genotype a better competitor than another? Antonie Leeuwenhoek 1998; 73: 35–47.

