## Supplemental Methods for "Adaptation to overflow metabolism by mutations that impair tRNA modification in experimentally evolved bacteria"

**Supplemental Text**

**Supplemental Materials and Methods**

All primers for cloning, mutagenesis, and tRNA production were obtained from Integrated DNA

Technologies (www.idtdna.com); sequences are provided in Supplementary Table 1.

The experimental GMM medium was made as (36.8 mM NaPO_4_ dibasic, 19.2 mM KPO_4_ monobasic, 0.79mM CaCl_2_, 3.16mM MgSO_4_), unintentionally with a lower buffering capacity. The original evolution experiment was conducted at the University of New Hampshire (UNH, Durham NH) with media solutions in deionized water, and most subsequent analyses were conducted at the University of Pittsburgh (Pittsburgh PA). Because phenotypes of mutant strains varied in reproducibility between laboratories, we sent water samples from both labs to Environmental Service Labs (Pittsburgh PA) to compare levels of trace elements. Levels of Zn, Fe, Na, and K differed significantly between samples, so all experiments reported here were conducted in Milli-Q filtered water from Pittsburgh supplemented with trace elements designed to mimic the UNH water used in the initial selection experiment (Supplementary Table 2).

**Experimental evolution and mutant collection.** Overnight cultures of HI2424^lac^ and HI2424^lac-^ genotypes were grown in T-Soy broth from freezer stocks and then sub-cultured into GMM as independent replicates to acclimate to the selection environment for 24h. Selection experiments were initiated by mixing HI2424^lac^ and HI2424^lac-^ at a 1:1 ratio in 5 mL GMM in 18x150mm test tubes, which were incubated at 37 °C in a roller drum at 30 rpm. Populations were propagated by daily 1:100 dilutions from ~10^8^/mL to ~10^6^/mL for six days (6.6 generations/day) and then 1:10,000 dilutions to 10^5^/mL for six additional days (10 generations/day) into new GMM. Plating occurred on on tryptic soy agar plates [30 g/L tryptic soy broth powder, 15 g/L agar and 60 μg/mL X-gal] to determine the frequency of the Lac marker. Growth curves were conducted using cultures founded from single colonies on agar plates in tryptic soy broth overnight, diluted 1:100 into 1% GMM (base medium), and preconditioned for 24 hours. All liquid cultures were grown in 5 mL at 37 °C on a roller drum. To start each competition experiment, pre-conditioned cultures were serially diluted 1:10,000 to match the selective bottleneck size.

**Whole genome sequencing and variant detection.** DNA was extracted from each evolved strain with putative beneficial mutations using the Wizard Genomic DNA Purification Kit (Promega Inc.). Sequencing libraries were prepared using a modified Illumina Nextera protocol and analyzed on an Illumina NextSeq 500 to a minimum average of 30x coverage [1]. Raw reads were processed using Trimmomatic to remove Nextera PE adapter sequences [2]. Processed reads were mapped to the *B. cenocepacia* HI2424 reference genome and mutations identified using the variant calling program breseq v0.31 or later [3].

**Growth curve methods and statistical analyses**: Growth curve assays were conducted in 96-well plates with replicates randomized to eliminate position-specific artifacts. Growth was monitored at 37°C with continuous shaking over 48 hours in a Tecan microtiter plate reader with optical density measurements at 600 nm (OD_600_) taken every 10 minutes. At least three independent experiments were performed with three biological replicates (nine total curves per strain) for each supplement. Analyses were performed in R (v4.0.5)[4] with several packages: area under the curve (AUC) was calculated with DescTools [5]; (v0.99.41); lag time and maximum log-linear growth rate v_max_ with growthrates [6] (v0.8.2); two-way ANOVA with base R; estimated marginal means and post-hoc Šidák-corrected pairwise means testing with cut off of 0.05 with emmeans [7] (v1.6.0); and compact letter displays assigned with multcomp [8](v1.4-17).

To evaluate relationships between cell density and pH trajectories, replicate 5mL cultures of WT and *tilS* N274Y were grown on a roller drum at 37 °C and destructively sampled to quantify OD_600_ and pH with a microtiter plate reader (Tecan Spark) and pH electrode (Fisherbrand accuTupH). Comparisons of growth in modified and CSH M9 media bases were conducted with samples every two hours from 12 to 36 hours, using two replicates staggered 12 hours apart for a total of 8 biological replicates per strain and 4 replicates per time point. Effects of carbon source were conducted by measurements at 0, 24, and 48 hours with three biological replicates.

**Preparation and radiolabeling of tRNA substrate.** *B. cenocepacia* tRNA^Ile2^(CAU) and tRNA^Met^(CAU) transcripts were synthesized as previously described [9] using sequences obtained from the Genomic tRNA database [10]. Double-stranded tDNA was constructed from overlapping oligonucleotides. Following T7 RNA polymerase run-off transcription and treatment with RNase-free DNase I (Thermo Scientific) the transcription product was recovered by ethanol precipitation. Following purification on a 10% denaturing polyacrylamide gel, tRNA was eluted from crushed gel fragments in 500 mM ammonium acetate (pH 5.3) with 1 mM EDTA. The tRNA was again ethanol precipitated and resuspended in 10 mM Tris-HCl (pH 7.5) and 1 mM EDTA. Transcripts were ^32^P-radiolabled at the 3´-internucleotide linkage with [α-^32^P] ATP using tRNA nucleotidyltransferase as described [11, 12].

**Enzyme cloning and expression.** *B. cenocepacia* HI2424 *metG* and *tilS* sequences were retrieved from the *Burkholderia* Genome Database [13]. Chromosomal DNA (cDNA) was isolated from cells using a Qiagen Qtip kit (Qiagen); genes of interest were amplified from cDNA and sub-cloned into a pET-28a overexpression vector (Novagen) between the NcoI/HindIII sites for *metG* and the NheI/HindIII sites for *tilS*. The *metG* and *tilS* genes were incorporated with His_6_-affinity tags at the C-terminal and N-terminal ends, respectively. TilS variants were generated from the initial construct using QuikChange mutagenesis (Agilent).

His-tagged wild-type and variant proteins were purified from *E. coli* Rosetta II (DE3) cells (Invitrogen) grown at 37 °C in LB with 50 μg/mL kanamycin (LB-Kan). Expression was induced with 1 mM IPTG (Fisher Scientific) at OD_600_ between 0.4 and 0.6. Cells were harvested 4 hours post induction, resuspended in buffer A (20 mM Tris-HCl [pH 8.0], 150 mM NaCl, 10 mM imidazole), and disrupted by sonication. Enzymes were purified on 5 mL Ni-NTA columns (GE Healthcare) pre-equilibrated with buffer A with elution using Buffer B (20 mM Tris-HCl [pH 8.0], 150 mM NaCl, 500 mM imidazole). His-tagged MetRS and TilS enzymes were recovered to greater than 90% homogeneity as determined by SDS-PAGE and were stored at -20 °C in 40 mM Tris-HCl (pH 8.0), 200 mM NaCl, 20 mM MgCl_2_, 20 mM KCl, and 40% glycerol. Concentrations were determined by UV absorbance (Thermo Scientific, NanoDrop 2000c). No modifications to the purification protocol were required for TilS variants.

**Lysidinylation by TilS.** *In vitro* lysidine synthesis activity was monitored as previously described [14, 15]. tRNA^Ile2^ (CAU) was annealed prior to analysis by heating to 80 °C for 2 minutes in 20 mM HEPES (pH 7.8), followed by slow cooling to room temperature. During cooling, MgCl_2_ was added to a final concentration of 10 mM. Lysidinylation was initiated by the addition of BcTilS (0.5 μM final concentration) to the reaction containing 100 mM Tris-HCl (pH 7.5), 5 mM DTT, 10 mM MgCl_2_, 10 mM KCl, 2 mM ATP, and 25 μCi ^3^H-lysine (Perkin Elmer) in the presence of 1 mM lysine and 2 μM tRNA transcript. The reaction proceeded at room temperature, and aliquots were quenched on Whatman filters pre-soaked in 5% trichloroacetic acid (TCA) and washed four times for 15 minutes each in 5% TCA. Lysidinylation activity of wild-type BcTilS and variants was determined from initial rates of reaction, and activities are reported relative to the wild-type value.

**Electrophoretic mobility shift assay.** tRNA^Ile2^ was annealed as for activity assays above. Each protein solution (220 nM-39.0 μM) was incubated with 3.5 nM ^32^P-labeled tRNA^Ile2^ at 37 °C for 30 minutes in 100 mM Tris-HCl (pH 7.8), 5 mM DTT, 10 mM MgCl_2_, 10 mM KCl, and 10% glycerol (10 μL total volume). The complex was separated from unbound tRNA on a native 10% polyacrylamide gel using a native gel running buffer (135 mM Tris pH 8.3, 960 mM glycine, and 5 mM EDTA) at 200 V for 20 minutes. The gel was dried, wrapped in plastic, and exposed overnight to a phosphorimager screen. The screen was analyzed using an Amersham Biosciences Storm 840 phosphorimager and ImageQuant 5.0 software. The fraction of bound tRNA was plotted against the concentration of TilS protein, and the data were fitted to a sigmoidal binding curve using GraphPad Prism v7.00 software to identify β_Max_, K_d_, and AUC for each protein sample.

**Circular dichroism**. Circular dichroism (CD) was conducted in 20 mM Tris-HCl (pH 7.8). Counter ions and salts were removed prior to CD by overnight dialysis. Protein concentrations varied depending on the TilS variant (13 μM wild-type, 6.2 μM A244T, 4.6 μM N274Y; 6.6 μM P421L, or 1.0 μM N445K). Molar ellipticity was monitored across 190-250 nm in an Aviv CD spectrometer, and percent α-helix was calculated from the absorbance at 222 nm [16]. Three technical replicates were averaged for each of three biological replicates. The average molar ellipticity was plotted against the wavelength using Prism software.

**Northern blot.** Total RNA isolated from ancestral and evolved strains of *B. cenocepacia* was isolated by Trizol extraction, ethanol precipitated and resuspended in Optima LC/MS grade water. Following extraction, a 10 μg sample was combined with 2X loading buffer and heat denatured at 80 °C, then separated on a 10% Urea-PAGE gel at 165 V for 1 hour. The RNA was transferred to a Biodyne B Nylon Membrane by electrophoresis in 1X transfer buffer (8 mM Na_2_HPO_4_ and 6 mM Na_3_C_6_H_5_O_7_) for 2 hours at 250 mA then an additional 2 hours at 350 mA in a 4 °C cold room. After transfer, the membrane was crosslinked for 1 min at 254 nm and 1,200 mJ twice using a UVP Hybrilinker Oven (Analytik Jena AG). The membrane was incubated in a warmed hybridization bottle with 8 mL ULTRAhyb-Oligo Hybridization buffer for 2 hours at 42 °C. After pre-hybridization, 1000 pmol of Cy5 labeled oligo, specific for either BctRNA^Met^ or BctRNA^Ile2^, was added and allowed to incubate overnight in a hybridization oven with rotation. The following day the membrane was washed twice with low-stringency buffer (2X SSC, 0.1% SDS) for 5 minutes each followed by two high-stringency (0.1X SSC, 0.1% SDS) washes for 15 minutes each. The membrane was visualized on an Amersham AI600 imager at 630 nm. Data analysis was performed using ImageQuant 7.0; BctRNA^Ile2^ was normalized to the BctRNA^Met^ intensity which served as an internal standard. The resulting data was used to compare relative lysidine abundance to the relative tRNA availability.

**Cellular lysidine content.** Cellular lysidine levels were determined by LC-MS as described [17]. Total RNA was isolated from ancestral and evolved strains of *B. cenocepacia,* from which 40 μg samples were folded in the presence of 10 mM MgCl_2_. Samples were brought up to 150 μL with Optima LC/MS grade water and then digested to individual nucleosides using P1 nuclease (SigmaAldrich) in 10 mM ammonium acetate (pH 5.3) at 50 °C for 2 hours. Reactions were quenched with 20 μL 0.1 M ammonium bicarbonate followed by the addition of 20 μL CutSmart buffer and 10 units of shrimp alkaline phosphatase (New England Biolabs). Phosphatase digestions proceeded at 37 °C for 2 h prior to inactivation at 65 °C for 5 min. Samples were centrifuged at 16,873 x g for 10 min; 196 μL of the supernatant was collected and 5 μL methanol:formic acid (80:4) was added prior to high performance liquid chromatography (HPLC) coupled electrospray ionization mass spectrometry (ESI-MS).

tRNA-derived nucleosides were separated by HPLC using a C-18 column (Polaris 3 100 x 4.6mm) and analyzed by a coupled mass spectrometer (Thermo LTQ Orbitrap XL). Solvent A was Optima LC-MS grade water with 0.1% formic acid, and solvent B was Optima LC-MS grade methanol with 0.1% formic acid. Separation was achieved at a flow rate of 0.3 mL/min with 15 μL injections by a gradient of 2% over 4 min, 2-100% over 21 min, 100% solvent B for 8 min followed by 100-2% over 6 min. The mass spectrum was collected in positive mode under the following conditions: voltage 4.01 kV, sheath gas flow rate 47, auxiliary gas flow rate 30, sweep gas flow rate 0.00, capillary voltage 2.00 V, capillary temperature 350.00 °C, tube lens voltage 49.89 V. *In vitro* lysidinylated tRNA was used as a standard. Lysidine in each sample was normalized to dihydrouridine as an internal standard.

**RNA sequencing and analysis of transcriptomes**. We followed the methods described previously [18] with modifications noted below. 0.5 mL of three independent preconditioned cultures were added to 50 mL of GMM in flasks and incubated at 100 rpm at 37 °C until reaching OD600 of 0.5. Each tube was centrifuged at max speed for 15 min at 25 °C and resuspended in cold PBS. 500uL of RNAprotect was added to the cell pellet followed by RNA extraction with Amresco Phenol Free RNA kits. The RNA was sequenced (1x75) on a NextSeq500 and the reads were pseudo-aligned to the HI2424 genome using Kallisto version 0.46 and then counted at 1000 bootstraps per sample [19]. Differential gene expression analysis was conducted using DESEQ2 [20] with scripts provided at <https://github.com/cdeitrick/rnaseq> and visualized using EnhancedVolcano [21]. The raw reads are available at NCBI Bioproject PRJNA895541.

**Supplemental References**

1.	Baym M, Kryazhimskiy S, Lieberman TD, Chung H, Desai MM, Kishony R. Inexpensive Multiplexed Library Preparation for Megabase-Sized Genomes. *PLoS ONE* 2015; **10**: e0128036.

2.	Bolger AM, Lohse M, Usadel B. Trimmomatic: a flexible trimmer for Illumina sequence data. *Bioinformatics* 2014; **30**: 2114–2120.

3.	Deatherage DE, Traverse CC, Wolf LN, Barrick JE. Detecting rare structural variation in evolving microbial populations from new sequence junctions using breseq. *Front Genet* 2015; **5**: 468.

4.	R Core Team. R: A language and environment	for statistical computing. 2020. R Foundation for Statistical Computing, Vienna, Austria.

5.	Signorell A, Aho K, Alfons A, Anderegg N, Aragon T, Arachchige C, et al. DescTools: Tools for Descriptive Statistics. 2021.

6.	Petzoldt T. growthrates: Estimate Growth Rates from Experimental Data. 2020.

7.	Lenth RV, Buerkner P, Herve M, Love J, Miguez F, Riebl H, et al. emmeans: Estimated Marginal Means, aka Least-Squares Means. 2022.

8.	Hothorn T, Bretz F, Westfall P, Heiberger RM, Schuetzenmeister A, Scheibe S. multcomp: Simultaneous Inference in General Parametric Models. 2022.

9.	Sherlin LD, Bullock TL, Nissan TA, Perona JJ, Lariviere FJ, Uhlenbeck OC, et al. Chemical and enzymatic synthesis of tRNAs for high-throughput crystallization. *RNA* 2001; **7**: 1671–1678.

10.	Chan PP, Lowe TM. GtRNAdb 2.0: an expanded database of transfer RNA genes identified in complete and draft genomes. *Nucleic Acids Res* 2016; **44**: D184–D189.

11.	Wolfson AD, Uhlenbeck OC. Modulation of tRNA^Ala^ identity by inorganic pyrophosphatase. *Proc Natl Acad Sci* 2002; **99**: 5965–5970.

12.	Bullock TL, Uter N, Amar Nissan T, Perona JJ. Amino Acid Discrimination by a Class I Aminoacyl-tRNA Synthetase Specified by Negative Determinants. *J Mol Biol* 2003; **328**: 395–408.

13.	Winsor GL, Khaira B, Van Rossum T, Lo R, Whiteside MD, Brinkman FSL. The Burkholderia Genome Database: facilitating flexible queries and comparative analyses. *Bioinformatics* 2008; **24**: 2803–2804.

14.	Nakanishi K, Bonnefond L, Kimura S, Suzuki T, Ishitani R, Nureki O. Structural basis for translational fidelity ensured by transfer RNA lysidine synthetase. *Nature* 2009; **461**: 1144–8.

15.	Nakanishi K, Fukai S, Ikeuchi Y, Soma A, Sekine Y, Suzuki T, et al. Structural basis for lysidine formation by ATP pyrophosphatase accompanied by a lysine-specific loop and a tRNA-recognition domain. *Proc Natl Acad Sci* 2005; **102**: 7487–7492.

16.	Correa DHA, Ramos CHI. The use of circular dichroism spectroscopy to study protein folding, form and function. *Afr J Biochem Res* 2009; **3**: 164–173.

17.	Edwards AM, Addo MA, Dos Santos PC. tRNA Modifications as a Readout of S and Fe-S Metabolism. In: Dos Santos PC (ed). *Fe-S Proteins: Methods and Protocols*. 2021. Springer US, New York, NY, pp 137–154.

18.	Mhatre E, Snyder DJ, Sileo E, Turner CB, Buskirk SW, Fernandez NL, et al. One gene, multiple ecological strategies: A biofilm regulator is a capacitor for sustainable diversity. *Proc Natl Acad Sci* 2020; **117**: 21647–21657.

19.	Bray NL, Pimentel H, Melsted P, Pachter L. Near-optimal probabilistic RNA-seq quantification. *Nat Biotechnol* 2016; **34**: 525–527.

20.	Love MI, Huber W, Anders S. Moderated estimation of fold change and dispersion for RNA-seq data with DESeq2. *Genome Biol* 2014; **15**: 550.

21.	Blighe K. EnhancedVolcano: publication-ready volcano plots with enhanced colouring and labeling. 2022.
